# ThermiQuant™ AquaStream: A portable instrument for quantitative colorimetric isothermal nucleic acid amplification reactions in paper and tube formats

**DOI:** 10.64898/2026.01.15.699357

**Authors:** Bibek Raut, Gopal Palla, Jiangshan Wang, Skyler Campbell, Virendra Kumar, Kyungyeon Ra, Mohit S. Verma

**Author notes:** Corresponding author (MV).

## Abstract

Microfluidic paper-based analytical devices (µPADs) are an attractive format for colorimetric nucleic acid amplification tests (NAATs) because they enable low-cost, portable diagnostics in resource-limited settings. However, researchers often optimize colorimetric assays in liquid reactions in tubes before translating them to µPADs. Since both formats require separate instruments for incubation and real-time sensing, direct comparison of reactions between the two formats is difficult. To address these cross-platform limitations, we developed ThermiQuant™ AquaStream, a portable benchtop device (15 × 20 × 16 cm, ∼5 kg; cost: USD 327) that supports seamless colorimetric loop-mediated isothermal amplification (LAMP) reactions in both µPADs and tubes under a common workflow. The system enables real-time reaction tracking (every 30 seconds) through onboard image processing, precise isothermal control (±0.5 °C) using a repurposed consumer-grade *sous-vide* heater, and medium-throughput (24 tubes or 42 µPADs). Testing with synthetic SARS-CoV-2 *orf7ab* DNA fragments demonstrated a limit of detection corresponding to a 95% probability of detection (LOD95) of 110 copies per reaction in tube (22 copies/µL) and 39 copies per reaction in µPADs (5 copies/µL), estimated using probit regression. In both formats the limit of quantification (LOQ), defined as the lowest concentration yielding a coefficient of variation (CV) of Tq ≤ 10%, was 250 copies/reaction resulting in a strong linear (R^2^ = 0.98 & R^2^ = 0.96 respectively for tube and µPADs) standard calibration curves. ThermiQuant™ AquaStream provides an affordable and versatile benchtop platform capable of supporting both tube- and µPAD-based colorimetric LAMP assays, serving as a proof-of-concept research tool for assay development and molecular diagnostics in One Health settings such as clinics, farms, and field environments.

## Introduction

Microfluidic paper-based analytical devices (µPADs) represent a promising format for colorimetric nucleic acid amplification tests (NAATs) because they provide affordable, portable solutions for diagnostics in settings with limited laboratory infrastructure [1–4]. However, researchers typically optimize assays in liquid reactions in tubes before translating them to µPADs, and both formats require separate instruments for incubation and real-time sensing [3,5–8]. This fragmented setup can make it inconvenient to evaluate assays across formats during development. To address this limitation, we developed ThermiQuant™ AquaStream (Fig 1), a benchtop heater-imager that supports both tube- and µPAD-based colorimetric LAMP reactions within a single instrument platform.

**Fig 1.**
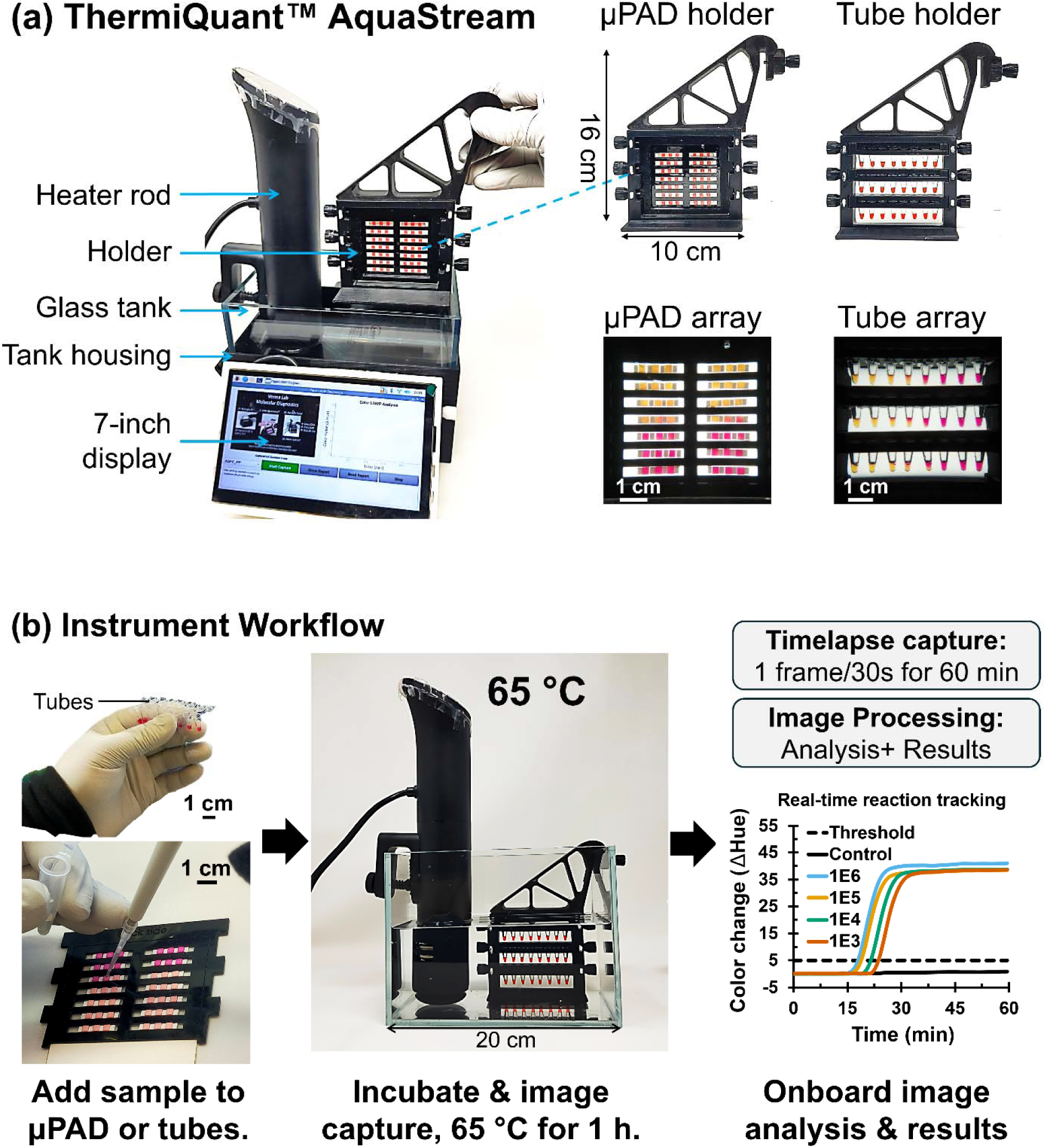
Design and workflow of the ThermiQuant™ AquaStream platform for dual-format (µPAD and tube) colorimetric NAATs. (A) Exploded view of the instrument with interchangeable holders for either 24 tubes or a 42-zone µPAD cartridge. (B) Typical experimental workflow: samples are loaded into tubes or a µPAD cartridge, incubated at 65 °C in the water bath for 1 hour with continuous imaging at 30 sec intervals, followed by automated hue-based color analysis and result reporting.

Colorimetric isothermal NAATs such as loop-mediated isothermal amplification (LAMP) [1,9] are practical alternatives to polymerase chain reaction (PCR) because they operate at a constant temperature and provide visible readouts [4,10]. These visual signals arise from chemical changes during amplification: either a drop in pH as protons accumulate, shifting dyes such as phenol red from pink-red to yellow, or the decrease in free Mg^2+^ which alters the color of metal ion-sensitive dyes such as hydroxynaphthol blue (violet to blue) or calcein (orange to green) [2,4,10,11]. However, uneven color development often prevents observers from easily distinguishing positive from negative reactions by the naked eye [3,12]. This challenge is heightened on µPADs, where color formation near the limit of detection (LOD) is often weak or spatially uneven [3,5,6]. Prior work with lateral flow and µPAD-based assays has shown that such subtle or irregular signals frequently lead to incorrect user interpretations, especially for borderline positives [13]. Consistent with these observations, our µPAD-LAMP pilot study also revealed notable misclassification due to ambiguous color outcomes [7]. Together, these findings highlight the limitations of relying on unaided visual judgment and underscore the need for devices that can reliably capture and analyze colorimetric signals to support accurate diagnostic decisions.

Equally important, researchers typically follow a staged workflow for µPAD-LAMP assay development. They first screen primers in liquid reactions using fluorescent dyes or probes, then test selected primers in liquid colorimetric assays, and finally translate them to µPADs while systematically optimizing dye concentrations and reagent formulations at each stage [5,8,11,14–16]. As summarized in Table 1, devices reported over the past five years reflect this fragmentation: researchers use commercial thermocyclers for fluorescence-based liquid assays that support full automation, while they commonly utilize custom-built analyzers to evaluate liquid colorimetric assays [17,18] or µPADs [14,19,20]. A few exceptions (e.g. Ubiquitous NAAT (UbiNAAT) [21], Microfluidics Integrated LED-Photodiode (MILP) device [22], LAMP-CRISPR device [23]) have attempted fluorescence detection directly on µPADs using custom optics integrated within smartphones or microcontroller-based cameras. Collectively, this fragmentation forces assay developers to switch between multiple instruments, increasing costs and reducing reproducibility and cross-validity when translating assays from liquid colorimetric to µPADs.

**Table 1:** Comparison of µPAD and tube based colorimetric and fluorometric isothermal NAAT instruments published since. 2020. Acronyms: PC, personal computer; PCB, printed circuit board; PTC; positive temperature coefficient; µPAD, microfluidic paper-based analytical device; n/a, not available.

| Instruments | Reaction types | Heater type | Color sensing | Cost (USD) | Size | Onboard/ offboard analysis |
| --- | --- | --- | --- | --- | --- | --- |
| ThermiQuan™ AquaStream (this work) | μPAD & liquid tubes | 65 ± 0.5 °C using water bath | Camera (Hue) | \$327 | 20 × 15 × 16 cm, 5 kg | Onboard on Raspberry Pi 4B |
| Smartphone operated LAMP (SMART-LAMP) [28] | Liquid tubes | 65 ± 0.9 °C using resistive heater | TCS3472 color sensor | €300 (~\$350) | 12.5 × 7.8 × 8.8 cm, 0.85 kg | Offboard on smartphone |
| Quantitative colorimetric LAMP (qcLAMP) device [18] | Liquid tubes | 63 ± 1-2 °C using PCB heater | Mini camera (RGB color differences) | n/a | 11 × 10 × 10 cm, 0.37 kg | On board on Raspberry Pi & on smartphone app |
| Integrated Gene box (i-Genbox) [17] | Custom liquid wells | 65 ± 1-2 °C using thin film heater | Smartphone (Hue) | n/a | 8 × 5 × 6 cm | On board on Smartphone |
| Human Papillomavirus (HPV) LAMP reader [24] | Custom liquid wells | 65 ± 0.5 °C using resistive heater | Color sensor | Consumable: \$5; Device: n/a | 0.29 × 0.21 × 10.6 cm | Onboard on a microprocessor |
| ThermiQuant™ MegaScan [20] | μPAD | 65 ± 0.5 °C using water bath | Scanner (Hue) | \$1,573 | 50 × 31 × 25 cm, 8 kg | Offboard on PC using Amplimetrics™ software |
| Field Applicable Rapid Microbial LAMP (FARM-LAMP) [19] | μPAD | 65 °C using water bath | Camera (Hue) | n/a | 16.4 x 13.5 x 19.3 cm | Offboard on PC using custom Python script |
| Ubiquitous NAAT (UbiNAAT) [29,30] | μPAD | 63 °C (but setpoint set to 71.5) using PCB heater | Smartphone with fluorescence optics | \$1,458 | ~13 x 3 x 8 cm | Offboard on PC using custom script |
| Paper LAMP-CRISPR device [31] | μPAD | 48 to 65 °C using thin film heater | Camera with fluorescence optics | n/a | 15 × 15 × 10 cm, <1.8 kg | Onboard using Raspberry Pi 3A+ |
| Microfluidics Integrated LED-Photodiode (MILP device) [22] | μPAD | 65 °C using thin film heater | Fluorescence optics | \$1,763 | Test kit: 4.5 × 2.0 × 2.0 cm, device: n/a | Onboard using Arduino |

For an integrated workflow, a next-generation colorimetric NAAT instrument must incorporate four key features (as quantified in Table 1): precise temperature control, accurate and reproducible color sensing, affordability and portability, and onboard analysis. Existing systems (Table 1) already demonstrate aspects of each. In terms of temperature control, portable printed circuit board (PCB)- and cartridge-based heaters achieve 0.5–2 °C accuracy [17,21,23,24], while benchtop water-bath systems offer even tighter regulation at ±0.5 °C [19,25,26]. For color sensing, flatbed scanners and cameras allow monitoring of reactions by tracking reaction-driven color changes [14,17,19,20]. Regarding cost and portability, analyzers such as Field Applicable Rapid Microbial LAMP (FARM-LAMP) [19], Integrated Gene box (i-Genbox) [17], and LAMP Assay Reader Instrument (LARI) [27] achieve shoebox-scale footprints (10–20 cm longest side, <3 kg) and prices far below commercial PCR instruments (>USD 10,000), typically in the USD 200–2000 range. Researchers have explored workflow automation through smartphone apps [17] and Python-based pipelines such as Amplimetrics™ [14], demonstrating both real-time and post hoc analysis. Recent advances demonstrate the technical feasibility of all four key features; however, an integrated instrument that combines them into a single compact platform capable of reproducible, side-by-side optimization across both liquid and µPAD assays is still lacking.

Learning from these developments (Table 1), we designed ThermiQuant™ AquaStream (Fig 1), a compact instrument that unifies tube- and µPAD-based colorimetric NAATs within a single workflow. AquaStream integrates the same four essential features identified above. Its name reflects these integrated features: “Therm” (heating), “i” (imaging), “Quant” (quantification), and “AquaStream” (water circulation). For temperature control, it delivers precise volumetric heating at 65 ± 0.5 °C using a repurposed consumer-grade *sous-vide* precision water heater. For color sensing, it incorporates backlit illumination optimized for both liquid tubes and µPADs in combination with a 16-megapixel autofocus camera (Fig 1 and 2). For cost and portability, AquaStream maintains a compact footprint (20 × 15 × 16 cm, 5 kg) with a bill of materials (BOM) component cost of USD 327. For workflow automation, it enables real-time analysis (every 30 seconds) onboard, with integrated image capture and processing across both formats. Collectively, these features enable continuous colorimetric LAMP kinetic monitoring and establish a unified, reproducible platform for assay optimization and translation across formats. Looking ahead, AquaStream has the potential to expand decentralized molecular testing, supporting One Health applications in the clinic, on farms, and outdoors.

**Fig 2.**
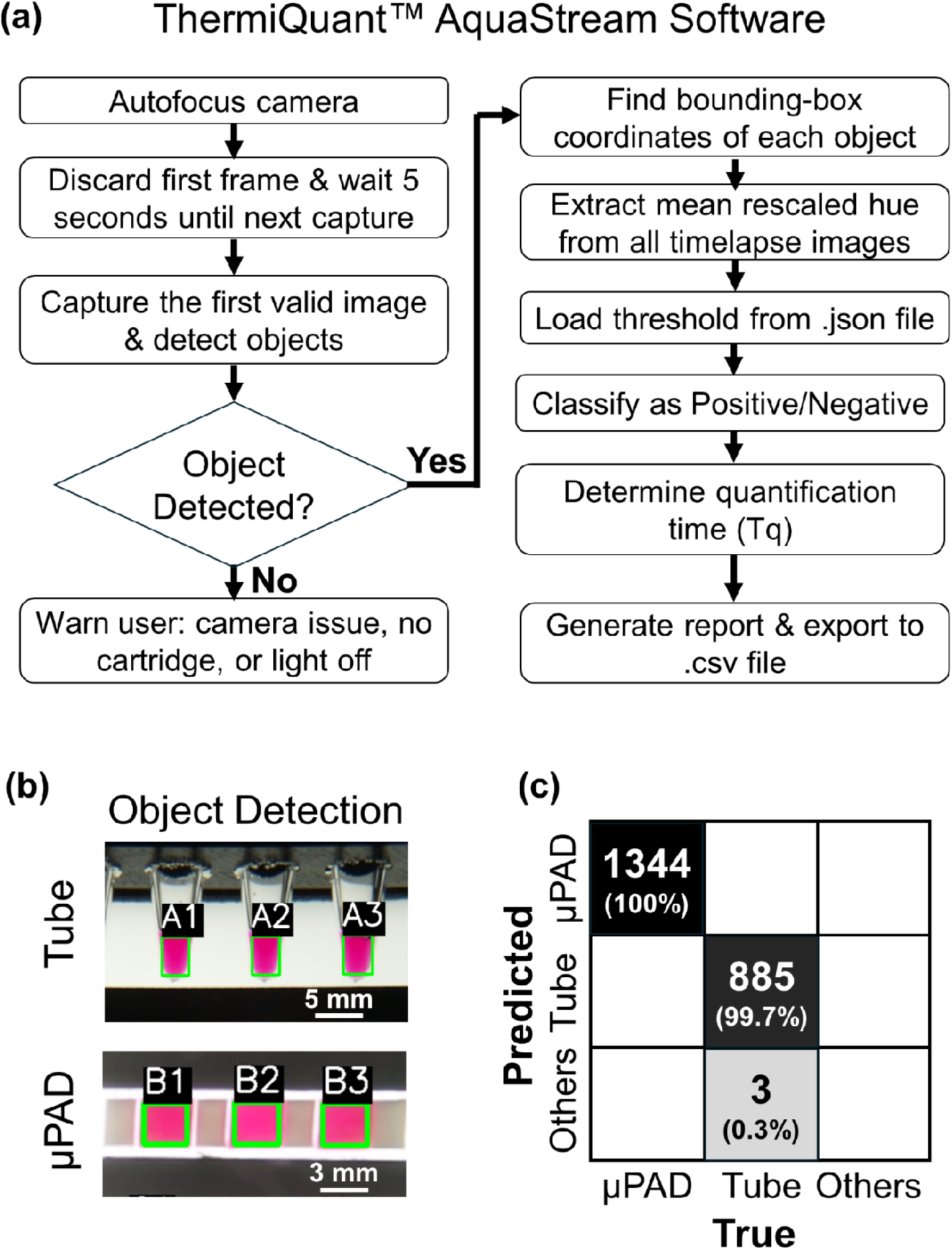
Automatic image analysis. (A) Software workflow for timelapse image capture and image processing. (B) YOLOv8n instance segmentation showing bounding box detection and automatic labeling of tubes and µPAD zones. (C) Confusion matrix summarizing detection accuracy for tubes and µPADs.

## Materials and Methods

### ThermiQuant™ AquaStream design, fabrication, and assembly

The ThermiQuant™ AquaStream has nine major components: (i) water heater rod (Anova Nano 3.0, ASIN: B0BQ93XGWC, “heater_rod.step”) (ii) 3D printed tank cover (“tank_lid_aquastream.step”), (iii) 3D printed cartridge holder (“cartridge_holder_base.step”, “cartridge_holder_handle.step”, “cartridge-clamp_paper-sensor.step”, “knob.step”, “tube_cartridge.step”, “tube_clamp”) (iv) glass water tank (Awxzom, ASIN: B0D3F36CTY, “glass_tank.step”), (v) backlight LED panel (Weilisi, WLS-PC02), (vi) camera (Arducam IMX519, SKU B0371, “camera_case.step”), (vii) Raspberry Pi 4B (4 GB RAM; Raspberry Pi Foundation, UK; “raspberry_pi4B_case.step”), (viii) 7-inch touch display monitor (Freenove, FNK0078B; “7-inch_display_holder.step”, “7-inch_display_holder-hook.step”), (ix) 3D printed tank box (“glass_tank_housing.step”). Exploded views of the assembled instrument are shown in Fig 1A.

All 3D models were designed using SolidWorks (Dassault Systèmes SolidWorks Corp., USA), and the corresponding design files are available in the Supporting Information (“Design_files” folder). The tank box and cover were 3D-printed with 15% infill density from polyethylene terephthalate glycol-modified (PETG) filament using a Bambulab X1 Carbon printer (Bambulab, China), while the cartridge holders were printed with 100% infill density from polycarbonate filament. Both PETG and polycarbonate filaments (Overture, USA) were used selectively: PETG was chosen for external housing due to its low warping and ease of printing, while polycarbonate was used for cartridge holders and inserts placed inside the water bath because it withstands prolonged (tested for > 6h) exposure to hot water (65 °C) without softening.

Instrument assembly is described in S1 Fig within S1 file, cartridge assembly in S2 Fig, with the complete bill of materials (BOM) provided in S1 Table. The instrument workflow overview is shown in Fig 1B and step-by-step user guide shown in S3 Fig.

### Water bath temperature characterization

To assess thermal equilibration and spatial uniformity, four thermocouples were positioned diagonally across an acrylic cartridge at 20 mm intervals (Fig 3A, top panel). The cartridge was then submerged in a water bath stabilized at 65 °C, and temperature readings were continuously collected for 30 min using a multichannel data logger (AZ Instruments K-type thermocouple recorder with 8G SD card, Amazon, USA). From these measurements, both equilibration time and spatial variability among the probes were determined.

**Fig 3.**
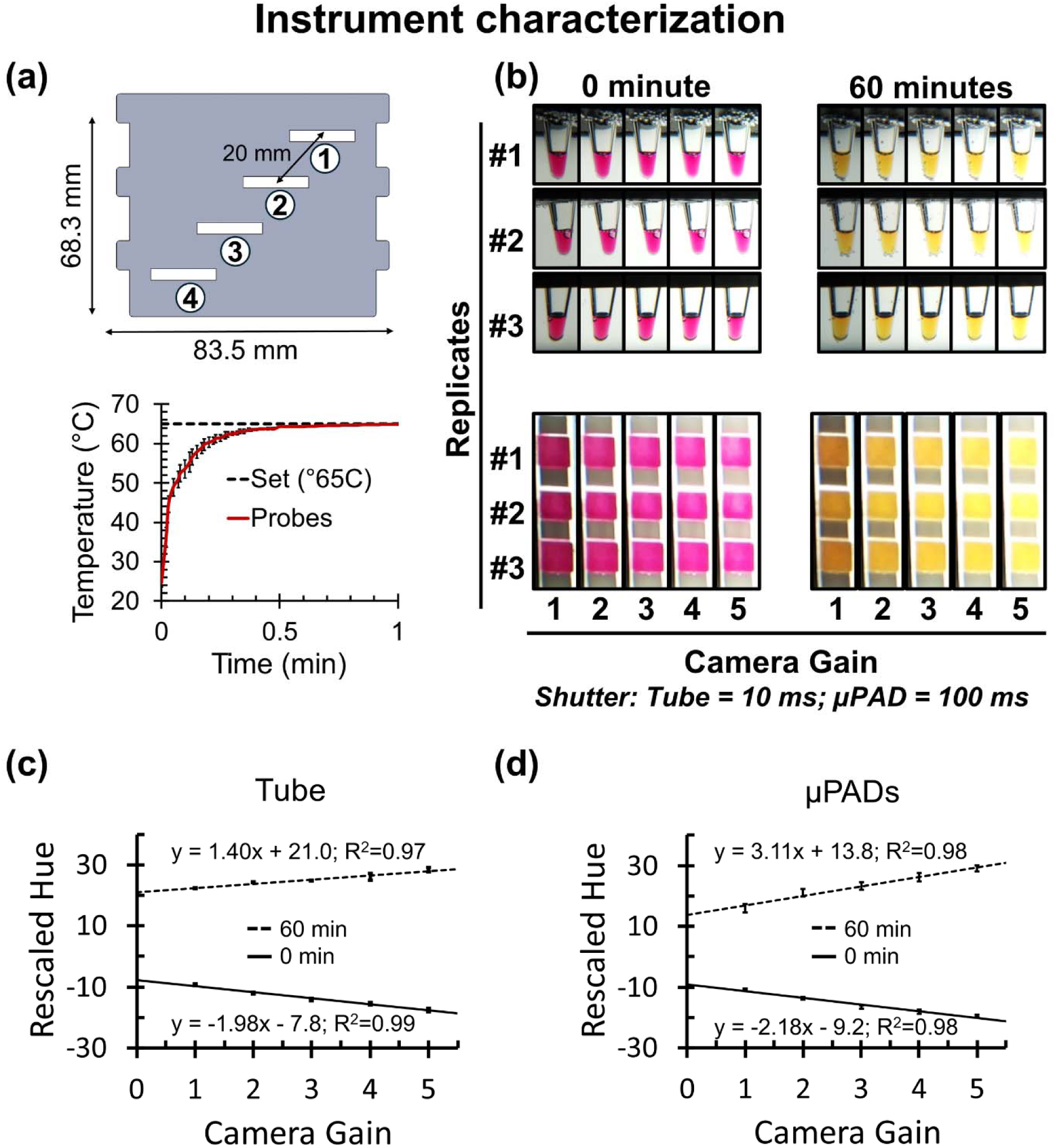
Instrument characterization. (A) Top panel shows thermocouple placement at four diagonal positions (1–4) across the cartridge, spaced 20 mm apart. Bottom panel show average temperature of four temperature probes and error bars represent standard deviation. Temperature equilibrated in less than 1 minute at 65 °C with spatial variation maintained within ±0.5 °C across probes. (B) Representative images of tube and µPAD reactions acquired at varying camera gain settings, with exposure times fixed at 10 ms (tubes) and 100 ms (µPADs). Images shown at 0 and 60 minutes illustrate negative and positive colorimetric endpoints of LAMP reaction across three replicates. (C) Raw hue values versus camera gain for tubes, showing strong linear correlations at both 0 min (R^2^ = 0.99) and 60 min (R^2^ = 0.97). (D) Corresponding hue versus gain plots for µPADs, also showing strong correlations (R^2^ = 0.98 at both 0 and 60 minutes).

### Synthetic *orf7ab* DNA preparation and copy number calibration

We used the procedure from our previous report [20]. Briefly, synthetic DNA corresponding to the SARS-CoV-2 *orf7ab* gene sequence (NCBI Reference Sequence: NC_045512.2; S2 Table) were obtained as 4 ng lyophilized pellets (IDT, USA). The pellets were rehydrated to a final stock volume of 40 μL using nuclease-free water (Fisher Scientific, 43-879-36). Primers and probes for digital PCR (dPCR) specific to the *orf7ab* target were obtained from previously published work [20] and are listed in S3 Table.

Each dPCR reaction was assembled in a total volume of 40 μL, containing 10 μL of 4× Probe PCR Master Mix (Qiagen, 250102; final concentration 1×), 4 μL of 10× primer–probe mix (final concentration 1×; 0.8 μM forward primer, 0.8 μM reverse primer, and 0.4 μM FAM-labeled probe; see S3 Table), 0.5 μL of EcoRI-HF restriction enzyme (New England Biolabs, R3101S), 20.5 μL of nuclease-free water, and 5 μL of DNA template. Reaction mixtures were loaded into a 26K 24-well Nanoplate (Qiagen, 250001) and analyzed on a QIAcuity One 5-plex dPCR system (Qiagen, 911021). The thermal cycling profile included an initial denaturation step at 95 °C for 2 min, followed by 40 cycles of 95 °C for 15 s, 55 °C for 15 s, and 60 °C for 30 s. Absolute quantification of the DNA copy number was performed using the QIAcuity Software Suite (Qiagen). The quantified DNA stock solution was stored at -80 °C until further use.

### Liquid LAMP experiment setup

The colorimetric LAMP liquid-phase assay was based on the formulation reported in our earlier study for paper LAMP [20,32], with minor modifications. In the current study, we used the same formulation for liquid and paper LAMP assays to allow comparison between the two formats. A 1,000 μL 2× LAMP mix was prepared by combining 100 μL KCl (1,000 mM; Sigma-Aldrich, P9541), 160 μL MgSO (100 mM; Sigma-Aldrich, M2773), 280 μL dNTP mix (10 mM; Fisher Scientific, FERR0182), 2.8 μL dUTP (100 mM; Fisher Scientific, FERR0133), 0.4 μL Antarctic Thermolabile UDG (1 U/μL; New England Biolabs, M0372S), 5.4 μL Bst 2.0 DNA polymerase (120 U/μL; New England Biolabs, M0537M), 20 μL phenol red (25 mM; Sigma-Aldrich, P3532), 100 μL Tween-20 (20%; Sigma-Aldrich, P9416), and 331.4 μL nuclease-free water (Fisher Scientific, 43-879-36).

For the preparation of 200 μL of the final LAMP master mix, 125 μL of the 2× LAMP mix was combined with 25 μL of 10× primer mix (16 μM FIP/BIP, 2 μM F3/B3, and 4 μM LF/LB; final concentrations: 1.6 μM FIP/BIP, 0.2 μM F3/B3, 0.4 μM LF/LB; see S4 Table), 0.67 μL Bst 2.0 DNA polymerase, 1 μL betaine (5 M; Sigma-Aldrich, B03005VL), 3.13 μL bovine serum albumin (BSA) (40 mg/mL; Sigma-Aldrich, A2153), 36.0 μL trehalose (1.75 M; Thermo Scientific Chemicals, 182550250), and 9.2 μL nuclease-free water. The pH of the final master mix was adjusted to ∼7.9 with 0.1 M KOH, resulting in a uniform, red-colored solution.

The synthetic *orf7ab* DNA, quantified by dPCR, was serially diluted to generate target concentrations of 10^6^, 10^5^, 10^4^, 10^3^, 10^2^, 10, and 1 copy per reaction for the broad-range LOD experiment. A narrower dilution set: 10^3^, 5×10^2^, 2.5×10^2^, 10^2^, 50, 25, 10, and 1 copy per reaction was used for narrow range LOD evaluation. Each dilution was prepared with three technical replicates per concentration. Each LAMP reaction contained 20 μL of the colorimetric final LAMP master mix and 5 μL of the DNA template, resulting in a total reaction volume of 25 μL. For no-template controls (NTCs), an equivalent volume of nuclease-free water was substituted for the DNA template. The sealed reaction tubes were placed in a 3D-printed holder, inserted into the AquaStream device, and incubated at 65 °C for 60 min.

### µPAD LAMP experiment setup

µPAD strip sensors were fabricated following a previously published protocol [20,32,7,14]. Each strip consisted of three pairs of alternating 0.83 mm-thick Grade 222 chromatography paper pads (3 × 3 mm) and 0.5 mm-thick polystyrene spacers (2 × 3 mm), assembled on a 0.076 mm (3 mil) double-sided adhesive–layered MELINEX® polyester backer. Polystyrene spacers were used to create fluidic barriers between adjacent µPADs.

A custom cartridge was fabricated from 1.5 mm-thick acrylic sheets using a CO_2_ laser cutter (Glowforge Plus, 40 W). The laser cutter operated at 100% power and speed factor 160 for acrylic sheets and at 15% power and speed factor 150 for PCR sealing films. PCR plate seals (Fisher Scientific, AB-0558) were first attached to one face of the cartridge, after which µPAD strips were positioned inside using tweezers. A 7.5 μL aliquot of the final master mix (see Methods: “Liquid LAMP experiment setup”) was then pipetted onto each µPAD and allowed to dry inside a PCR workstation (Mystaire, MY-PCR32) for 2 h.

dPCR quantified *orf7ab* DNA stocks were serially diluted to concentrations ranging across 10^6^, 10^5^, 10^4^, 10^3^, 5×10^2^, 2.5×10^2^, 10^2^, 50, 25, 10, and 1 copy per reaction. For each dilution, 7.5 μL of the DNA solution was pipetted onto the µPAD housed inside the cartridge. One strip was used per concentration group, and since each strip contained three µPADs, each group included three technical replicates. NTCs were also included, with nine in run 1 and five in run 2. The smaller number of NTCs in Run 2 resulted from µPAD damage, while the larger number in Run 1 was intentional to increase control replicates relative to dilution triplicates. After sample pipetting, the cartridges were sealed with PCR tape to prevent evaporation and leakage. S4 Fig illustrates the pipetting of diluted DNA samples into reagent-preloaded µPADs, followed by cartridge sealing with PCR tape.

### Image acquisition, processing, and analysis workflow

After placing the tubes or µPADs containing holders into a 65 °C preheated water bath and launching the python script onboard the Raspberry Pi 4B, the system captured time-lapse images for downstream analysis. Three experimental workflows were implemented. In the first workflow, images were captured using a custom Python script which collected frames every 30 s across gain settings 1-5, with 5 s intervals between gains (supporting information, S3 file, “Software\01_Capture_timelapse” folder; Python files: “paper_LAMP.py” or “liquid_LAMP.py”). In the second workflow, the time-lapse images were transferred and processed through an updated version of our previously published Amplimetrics™ software (supporting information, “Software\02_Amplimetrics-V1.2” folder; Python files: “Amplimetrics_V1.2.py”) [20]. The Amplimetrics™ V1.2 software was mainly updated with a YOLOv8-based instance segmentation model trained on a new dataset incorporating both µPADs and tubes (Fig 2B-C) [33]. The software automatically detected the regions of interest, extracted average hue values, and generated hue-versus-time plots. This workflow was used to identify the optimal camera gain, backlight intensity, and camera exposure duration settings. Further, the hue-time data for each sample was processed again in Amplimetrics™, where net hue change was obtained by subtracting all subsequent hue from their initial hue value. The positivity threshold was set as the upper bound of the 95% bootstrap confidence interval of the mean net hue change of the NTCs, estimated from 100,000 bootstrap resamples. For positive reactions, the quantification time (Tq) was determined as the point at which the second derivative of the hue–time curve reached its first maximum, corresponding to the phase of maximum kinetic acceleration in the colorimetric LAMP reaction within the logarithmic region of the amplification curve. In the liquid colorimetric LAMP reaction, we applied a time-based cutoff in addition to the hue-based positivity threshold to reduce the high incidence (5/21 or 24%) of amplification in NTCs. The time threshold was calculated by bootstrapping the Tq values of NTCs that were positive (n = 5) and taking the lower bound of the 95% confidence interval of the bootstrap mean. Reactions with Tq values above this threshold were classified as late amplification events and therefore considered negative. This time cutoff was not applied to the paper-based colorimetric LAMP reaction, as the incidence of NTC amplification was low (1/15, 6.7%). Plotting Tq against the log_10_ of template concentration produced a linear calibration curve, enabling quantitative prediction of target concentrations in unknown positive samples.

In the third workflow, the positivity threshold value and calibration equation obtained from Workflow 2 were overwritten into the final AquaStream app as a JSON file within the main program. This allowed subsequent experiments for the same template type to be executed in batch mode, with the software automatically providing both binary classification (positive/negative) and concentration estimates for unknown samples at the end of each run. The overall instrument workflow for this configuration is shown in S3 Fig, while the corresponding software workflow is illustrated in Fig 2. The software was run on Raspberry Pi 4B single board computer and the source code is located in the supporting information (“Software\03_AquaStream_V1.0” folder; Python files: “AquaStream_software.py”).

### Determination of LOD95 and LOQ

The LOD95 was determined by probit regression analysis [34]. Binary amplification outcomes (positive or negative) obtained from two independent experiments were combined (n = 6 replicates per concentration) and modeled as a function of the log_10_-transformed target concentration. The LOD95 was defined as the concentration corresponding to a 95% predicted probability of detection. Because only six replicates per concentration were used, the resulting value represents a preliminary estimate intended to demonstrate that the instrument is capable of performing and analyzing such assays. A full analytical validation was beyond the scope of this study. Regulatory guidance (e.g., FDA recommendations) typically requires testing at least 20 replicates of positive and negative samples near the expected LOD and determining the concentration at which ≥19 of 20 replicates produce positive amplification [35].

The LOQ was was established based on the precision of Tq measurements. For each concentration equal to or above the LOD95, replicate Tq values (n = 6) were used to calculate the coefficient of variation (CV = 100 × standard deviation / mean). The LOQ was defined as the lowest concentration yielding CV ≤ 10%, with at least three numerical replicate values required for CV calculation [36]. Although 35% CV is commonly used for qPCR assays, previous studies of LAMP assays report typical CV values of 5–10% for acceptable repeatability across reactions and runs; therefore, a 10% CV threshold was selected in this study [37–40]. For calibration analysis, linear regression of Tq against log_10_-transformed concentration was performed using replicate-level measurements at concentrations equal to or above the LOQ. The 95% prediction interval of the regression model was calculated to describe the expected variability of individual measurements.

## Results

### ThermiQuant™ AquaStream is compact, low cost, and supports dual-format colorimetric loop-mediated isothermal amplification reaction with high throughput

We developed ThermiQuant™ AquaStream as a compact, low-cost, modular instrument for real-time analysis of colorimetric LAMP reactions. Fig 1A illustrates the instrument design, while Fig 1B outlines the user workflow. The device integrates a water bath to maintain a constant incubation temperature of 65 °C and an onboard camera to continuously record the colorimetric transitions of LAMP reactions every 30 seconds. With a simple switch of the modular cartridge/tube holder, users can swap between 0.2-mL Eppendorf tubes and µPAD cartridges, enabling real-time analysis in both formats on the same platform. The instrument measures 15 cm × 20 cm × 16 cm and weighs ∼5 kg when filled with 3 L of water, making it readily portable. AquaStream can be fabricated for under USD 327 (estimated BOM cost only) using 3D-printed components and consumer-grade electronics, making it comparable in cost to reported cost-effective portable modules listed in Table 1. In terms of throughput, the tube insert accommodates 24 reactions (three rows of eight 0.2-mL tubes), while the µPAD insert supports two columns of seven strips, each strip containing three µPADs, for a total of 42 reaction zones. Each µPAD zone reproducibly retained 7.5 µL of sample without leakage or crossflow into neighboring µPADs, enabling reliable independent reaction zone analysis due to excellent fluid retention by capillary forces within the µPADs. The system’s throughput exceeds that of most portable systems, which typically support only 8 tubes or 12 µPADs. The sole exception is our previously reported ThermiQuant™ MegaScan, which can process up to 160 µPADs in parallel but is limited to paper-based assays, costs nearly five times more, and is twice as heavy [20]. In contrast, AquaStream balances throughput, dual-format compatibility, portability, and affordability, uniquely supporting both µPAD and tube workflows within a single compact instrument. This dual-format capability is particularly useful for research and assay development workflows, where liquid LAMP reactions are often optimized before translation to µPAD-based formats.

### Onboard software atop ThermiQuant™ AquaStream automatically captures timelapse images, detects regions of interest, and provides real-time reaction analysis

Onboard software running on ThermiQuant™ AquaStream automatically captures time-lapse images, detects regions of interest, and performs real-time reaction analysis. We designed an automated software pipeline (SI, “Software” folder) that performs color zone detection and analysis every 30 seconds during the standard 60-minute experiment. Fig 2A illustrates the workflow, which begins once the user places the holder in the 65 °C water bath and presses “Start” on the onboard touchscreen.

The first task of the software is to detect the presence and position of tubes or µPADs. To automate this step, we trained a custom YOLOv8n instance segmentation model [33], which achieved 100% detection accuracy across 1,344 µPAD zones and 99.7% across 888 tubes (Figs 2B–C). This detection, powered by a Raspberry Pi 4B single-board computer, enables fully user-independent operation by automatically identifying and cropping regions of interest for downstream analysis.

For each reaction zone, the software calculates the average hue value across the entire segmented region to track the pink-to-yellow transitions characteristic of positive amplification. Hue values outside the expected phenol-red’s red-to-yellow range are excluded to minimize the influence of artifacts such as bubbles, uneven color development, or localized color variations. The resulting hue–time trajectory is then used to determine positive or negative reactions and to extract the quantification time (Tq).

Using calibration curves of Tq vs log_10_(DNA concentration), the system further estimated the concentration of unknown samples. At the 60-minute endpoint, AquaStream generated a summary report on the touchscreen within 30 seconds, eliminating manual interpretation and ensuring reproducible, quantitative analysis across both tube- and µPAD-based assays.

### Water bath maintains reaction temperature at 65 ± 0.5 °C

The water bath used in AquaStream provided stable isothermal conditions for LAMP amplification. While LAMP reactions can tolerate incubation between 60-65 °C, reproducible outcomes typically require stability within ±1 °C [19,26]. AquaStream met this requirement, maintaining 65 ± 0.5 °C throughout the 60 min run. Fig 3A (top) shows the experimental setup, where four temperature probes were positioned 20 mm apart along a diagonal of an 8.35 × 6.83 cm cartridge. Fig 3A (bottom) shows the average probe readings of four temperature probes with error bar indicating standard deviation, immediately after the cartridge was immersed in the 65 °C water bath from room temperature (∼23 °C). The internal temperatures when cartridge with probes were emerged in 65 °C preheated water bath, equilibrated from room temperature to 65 °C within 0.5 minutes and remained stable thereafter, with maximum deviations at equilibrium below 0.5 °C. Separately, the 3 L water bath required ∼10 minutes to reach setpoint when heated from room temperature using a consumer-grade 750 W, 110 V *sous vide* circulator, which ensured rapid warm-up and uniform heat distribution across the bath. These results confirm that AquaStream delivers precise, reproducible temperature control suitable for reliable LAMP reactions.

### Backlight affects hue measurement in colorimetric LAMP reactions

Front-illuminated imaging systems often suffer from shadows and uneven lighting, and in a water bath system these issues are further compounded by both reflection and refraction, which distorts illumination from the front. This limitation was evident in our earlier FARM-LAMP [19] platform, where additional image-correction steps had to be applied to compensate for these artifacts. To overcome these issues, AquaStream employs backlight illumination. While backlight illumination minimizes shadows and uneven lighting, it also introduces sensitivity to light intensity. Because varying the backlight required stabilization time between camera settings, we fixed the backlight at its lowest intensity and systematically adjusted the camera gain while holding exposure constant, in order to evaluate the effect of backlighting on colorimetric readings and identify optimal settings for both tubes and µPADs. Fig 3B shows images captured at different gain levels for colorimetric LAMP reactions with three technical replicates at 0 and 60 minutes, representing negative and positive endpoints. Corresponding plots of raw hue versus camera gain for tubes (Fig 3C) and µPADs (Fig 3D) revealed strong correlations (R^2^ = 0.97–0.99 for tubes, R^2^ = 0.98 for µPADs). The standard deviation of hue within technical replicates was consistently within ±2 hue units, confirming that the observed effects arose from gain adjustments rather than replicate variability. Interestingly, hue shifted in opposite directions: low gain pushed reds toward pink, while yellows shifted from faint or orange-tinged toward a clearer, more intense yellow at higher gain, widening the separation between negative and positive signals. This widening improved discrimination, but gain above 3 caused color washout, as visibly seen in Fig 3B. To maintain a clear red-to-yellow contrast while avoiding both underexposure and color washout, we fixed the operating parameters at Gain 2, with exposure times of 10 ms for tubes and 100 ms for µPADs. We note, however, that this choice was made for consistency across experiments rather than technical necessity; any gain setting between 1 and 5 supports reliable image analysis provided parameters are fixed across runs and instruments. Further optimization of gain or color modeling was not pursued in this study.

### Automated color analysis enables objective classification and establishes a 110-copies per reaction LOD95 in liquid LAMP assays

We evaluated AquaStream’s performance on liquid colorimetric LAMP reactions in tubes using two separate dilution studies with a synthetic DNA fragment corresponding to the SARS-CoV-2 *orf7ab* gene [7], designed to assess reproducibility of a single instrument across separate runs (Fig 4). While SARS-CoV-2 is naturally an RNA virus, a DNA surrogate was selected to provide a stable and well-controlled input template that avoids matrix-related variability while still maintaining clinical relevance. In previous work, we have demonstrated one-pot reverse-transcription LAMP reactions, where the RNA template undergoes reverse transcription to cDNA at the initiation of the reaction, after which the LAMP process amplifies the resulting DNA [7,20]. Because purified DNA diluted in nuclease-free water was used in this study, sample pH and buffering effects were minimal. For complex biological samples with stronger buffering capacity (e.g., blood, nasal swabs, or urine), dilution or preprocessing may be required to prevent interference with phenol-red based colorimetric detection, as demonstrated in our previous studies using complex matrices [7,8,14,20]. Each experiment included three technical replicates for every concentration along with NTCs as negative controls. The first experiment tested a broad range of input concentrations with wider spacing (10^6^ to 1 copy/reaction), while the second focused on a narrower range with finer spacing (10^3^ to 10 copies/reaction). Fig 4A-B shows cropped time-stamp images at 0, 20, 40, and 60 minutes for different concentrations. As amplification progressed, the reaction color shifted from pinkish red to yellow due to the accumulation of acidic by-products that lowered pH [10]. The color transition was often spatially heterogeneous, with intermediate states showing patches of red and yellow within the same tube, making reliable visual classification of positive and negative reactions difficult when assessed by the naked eye. In several reactions, the color change appeared to initiate near the bottom of the tube and gradually propagate upward over time. While the exact cause of this spatial pattern was not investigated in this study, it may be related to localized reaction kinetics and mass-transport effects within the tapered tube geometry, such as sedimentation of reaction components, temperature gradients, or diffusion-limited mixing during amplification. Because such intermediate and spatially heterogeneous states complicate visual interpretation, we quantified the reaction progression computationally by extracting the average hue value within each tube and plotting the net hue-versus-time trajectories for each concentration (Fig 4C). Positive reactions followed characteristic sigmoidal curves, whereas negatives remained flat or linear. We performed additional NTC amplification experiment with 15 technical replicates (S5A Fig). Using the net hue change at 60 min from all NTCs across three experimental runs (n = 21), we defined the positivity threshold as the upper bound of the 95% bootstrap confidence interval of the mean net hue change. The bootstrap distribution (S6A Fig) yielded a threshold value of 14.8 net hue unit.

**Fig 4.**
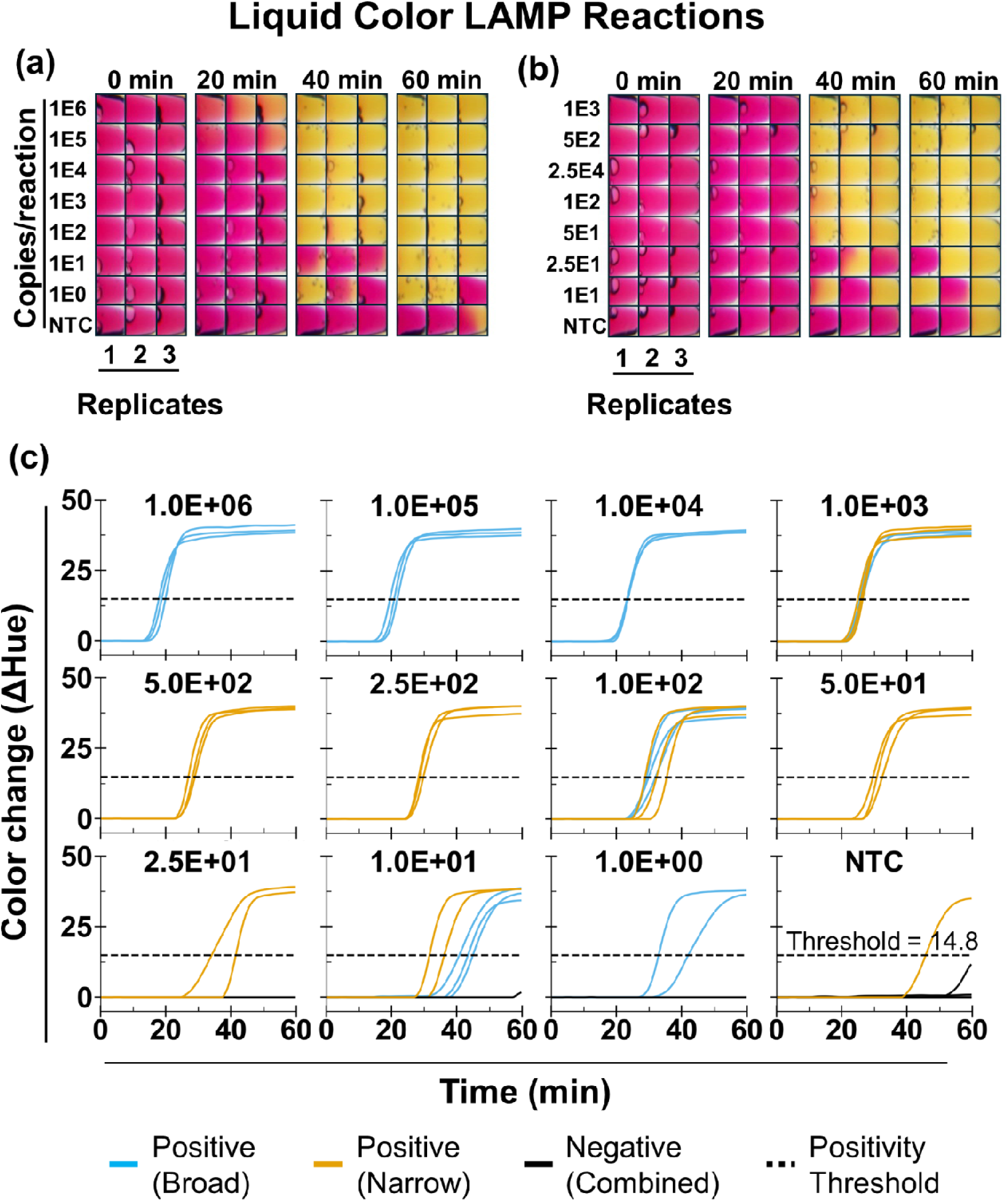
Analytical testing of colorimetric LAMP in tubes. (A) Broad-range dilution series from 10^6^ to 1 copies/reaction) with time-stamped cropped images (0, 20, 40, and 60 min) from tubes loaded with synthetic SARS-CoV-2 *orf7ab* DNA and no-template controls (NTC). Each concentration groups with 3 technical replicates. (B) Narrow-range dilution series from 10^3^ to 10 copies/reaction, also with three technical replicates per concentration. (C) Corresponding hue-versus-time trajectories, obtained from processed time-lapse images for each input concentration, showing sigmoidal shape for positive and linear or flat for negative. A positivity threshold of 14.8 net hue units was used for initial classification, followed by a quantification time (Tq) cutoff of 33.0 min (not shown) to determine final positives.

Using this positivity threshold, we observed false positives in 5 of 21 NTC reactions. We therefore used an additional Tq cutoff by bootstrapping the positive NTC Tq values and used lower bound of the 95% confidence interval of 33.0 min to define the Tq cutoff. This time cutoff reduced the number of false positives to 1 of 21 reactions (4.7%). Further, our LOD95 was 110 copies/reaction (22 copies/µL, S6B Fig).

We determined the LOQ to be 250 copies per reaction (50 copies/µL; S6C Fig), defined as the lowest tested concentration with a Tq coefficient of variation (CV) ≤ 10%. We selected the 10% CV threshold based on previous qLAMP studies, where CV values of 5–10% provide good quantification repeatability [37–40]. To further evaluate this LOQ, we performed linear regression of Tq versus log_10_-transformed standard concentrations, which showed strong linearity (R^2^ = 0.98; S6D Fig).

### Paper-based LAMP assays demonstrate reproducible detection across instruments

We next evaluated AquaStream’s performance with µPAD-based LAMP assays using the same synthetic DNA template as in the liquid experiments (Fig 5). Two independent but identical dilution studies were performed on separate AquaStream instruments of the same design, each including triplicate reactions per concentration as well as NTCs (9 on run 1 and 5 on run 2). Unlike the liquid experiments, which assessed repeatability across runs on a single instrument, these studies were designed to evaluate instrument-to-instrument variation. Figs 5A-B show representative time-stamp images at different input concentrations, with reactions transitioning from pinkish red to yellow as amplification progressed. As expected for porous paper substrates, color changes were spatially heterogeneous: some zones amplified rapidly while others showed delayed transitions despite identical input concentrations. This visual ambiguity was especially pronounced at lower concentrations (≤100 copies/reaction).

**Fig 5.**
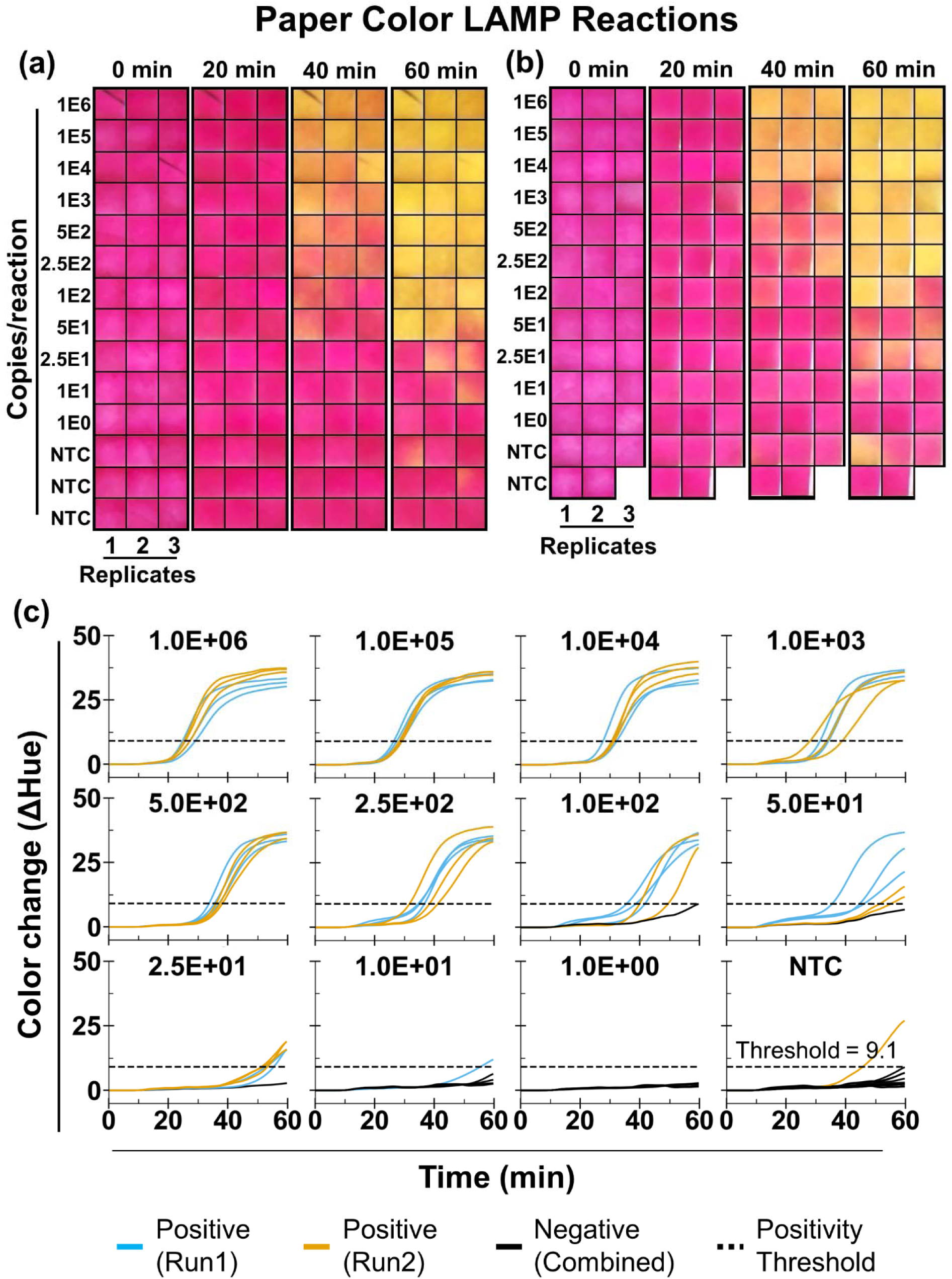
Analytical testing of colorimetric LAMP on µPADs. (A) Time-stamped cropped images (0, 20, 40, and 60 min) from µPADs loaded with serial dilutions of synthetic SARS-CoV-2 *orf7ab* DNA (10^6^-1 copies/reaction) with three technical replicates per concentration and no-template control (NTC) with 9 replicates. (B) Independent repeat of the same dilution series on a second AquaStream instrument of identical design, with 5 NTC replicates. (C) Corresponding hue-versus-time trajectories, obtained from processed time-lapse images for each input concentration. Reactions are classified as positive if they exceed the positivity threshold and as negative if they do not.

Automated hue-versus-time trajectories (Fig 5C) reliably distinguished positives from negatives despite this variability. Similar to the liquid LAMP analysis, we determined the positivity threshold as the upper bound of the 95% bootstrap confidence interval of the mean net hue change from n = 15 NTC reactions (S7A Fig). In both runs, amplification was consistently detected down to 50 copies/reaction, with all replicates crossing the positivity threshold (9.1 net hue unit). The LOD95 estimated using a probit model was 39 copies per reaction (5 copies/µL; S7B Fig). This value is comparable to the LOD95 previously reported for the ThermiQuant™ MegaScan instrument (34 copies per reaction) using the same SARS-CoV-2 assay on paper with a synthetic DNA target [20]. The assay time required to detect the LOD concentration was also similar, remaining under 60 minutes. Moreover, we observed only 1/15 false amplification on the NTC (6.7% false-positive rate). The LOQ was 250 copies per reaction (33 copies/µL), showing strong linearity (R^2^ = 0.96) and Tq CV ≤ 10%. The mean Tq bias between runs for concentrations at and above the LOQ was only 1.2 min with narrow 95% limit of agreement (-2.6 to 0.3 minute) between runs, indicating good inter-run repeatability (S9 Fig).

### Initial DNA concentration does not influence initial pH of LAMP reaction

To determine whether the addition of DNA could influence the initial color of the LAMP reaction in the low-buffer conditions, we evaluated both pH and image-derived hue values prior to amplification. We first divided nuclease-free water into two groups (n = 5 replicates per group) and measured their initial pH. In one group, synthetic DNA was added at 4E4 copies per µL (corresponding to 1E6 copies per reaction in a 25 µL tube reaction), whereas the other group remained as nuclease-free water only. A second pH measurement was then performed for both groups. The first and second pH in group 1 (positive sample) were 5.54±0.20 and 5.65±0.21 and in group 2 (negative samples) were 5.52 ±0.18 and 5.64 ±0.18. The change in pH (ΔpH), calculated as the second measurement minus the first measurement, was determined for each sample. As shown in S8A Fig, no statistically significant difference in ΔpH was observed between the DNA-spiked (positive) and non-spiked (negative) samples (p > 0.05), indicating that the addition of purified DNA does not measurably influence the initial pH of the reaction mixture. Additionally, to assess whether there was shift in initial reaction color, we analyzed the raw hue values at the initial time point for all tested DNA concentrations in liquid LAMP (Fig 4) and paper LAMP (Fig 5), including the NTC. As shown in S8 Figs B-C, we observed no statistically significant difference in hue between the NTC and DNA-spiked samples across the tested concentration range (1E6 to 1E0 copies per reaction) at time 0 min. In contrast, a fully amplified positive reaction (“PC60”) measured at the end point (60 min) produced the expected yellow color and showed a clear shift in hue compared with the NTC. These results indicate that adding purified synthetic DNA diluted in nuclease-free water does not introduce a measurable pH or color shift prior to amplification and that the observed colorimetric signal change arises primarily from the pH decrease generated during the LAMP amplification reaction itself.

### Cross-format quantification reveals liquid tube reaction faster than paper reaction by average 8.7 minutes

To demonstrate the instrument’s dual-format capability, we directly compared LAMP reactions performed in liquid and paper formats. Fig. 6A shows the standard calibration curves for both formats at and above the common LOQ of 250 copies per reaction, and Fig. 6B presents the Bland–Altman comparison of Tq values. On average, the Tq difference between paper and tube format was -8.7 min, indicating that liquid reactions amplified faster than paper-based reactions. The 95% limits of agreement were narrow, ranging from −7.9 to 9.4 min. These results indicate that liquid tube reactions show faster LAMP kinetics than paper-based reactions. Further investigation is required to understand the underlying mechanisms, such as potential adsorption effects or diffusion limitations in the paper matrix, which may influence amplification kinetics. Although this investigation is beyond the scope of the present study, the ThermiQuant™ AquaStream platform provides the capability to explore these effects in future work.

**Fig 6.**
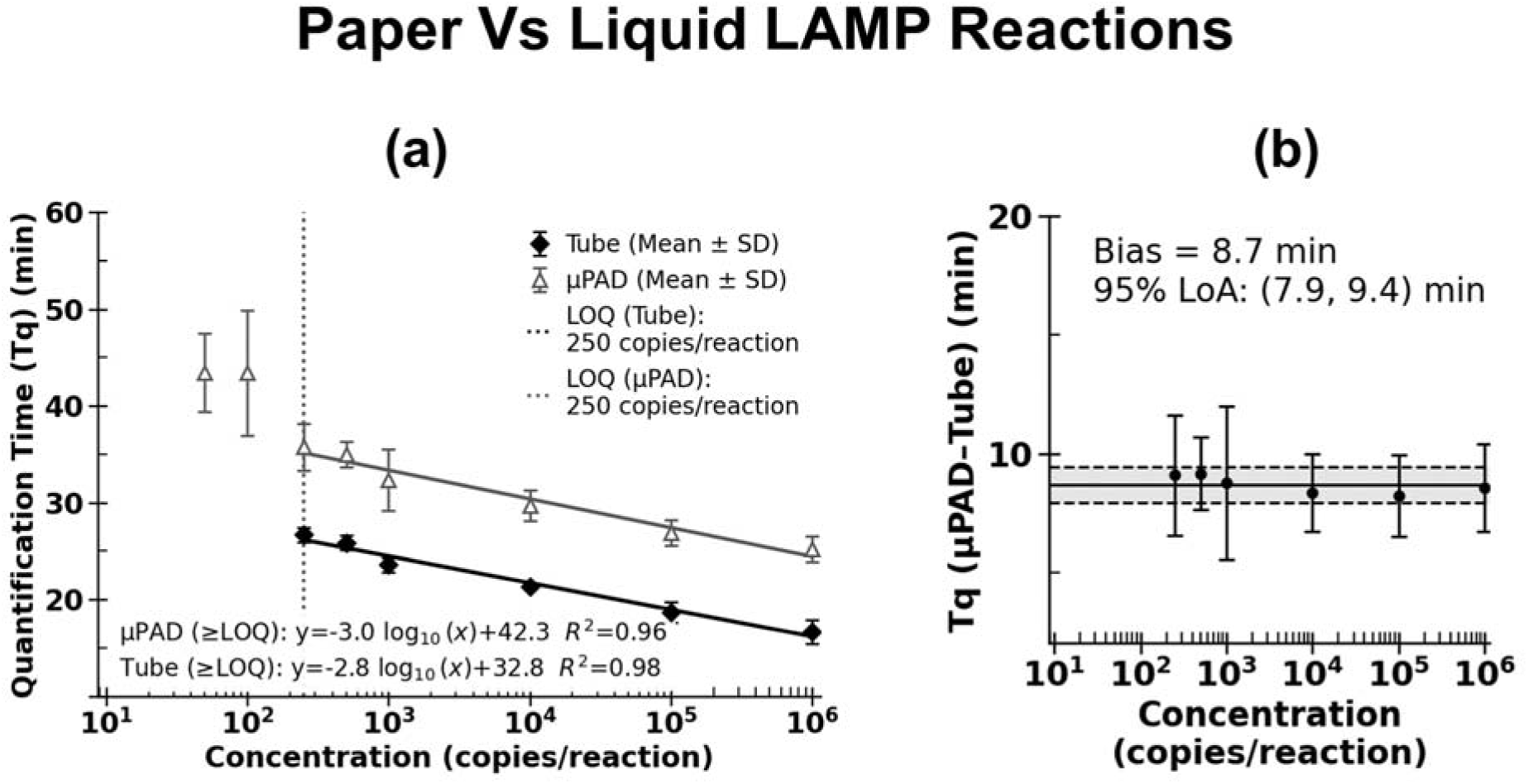
Cross-format quantification of colorimetric LAMP assays. (A) Quantification time (Tq) plotted against input concentration (log scale) for tube and µPAD assays at and above the LOD95, with linear regression performed only for concentrations at and above the LOQ. (B) Bland–Altman analysis comparing the Tq differences between liquid and paper LAMP.

## Discussion

We developed ThermiQuant™ AquaStream because, despite advances in colorimetric isothermal NAATs such as LAMP, no existing instrument enables parallel analysis of both µPAD- and tube-based reactions within a unified workflow. As summarized in Table 1, most instruments support only one format at a time, requiring separate setups for assay optimization and translation. AquaStream fills this gap by integrating four key features: precise temperature control, real-time imaging, portability and low cost, and automated data analysis.

For temperature control, we achieved stable isothermal heating at 65 ± 0.5 °C across the full cartridge, with the reaction core equilibrating to the water bath temperature in under a minute while maintaining a clear optical window for imaging (Fig 3A). This performance meets the ±1°C stability required for reproducible LAMP [41,42], matches the 0.3–0.5 °C precision of custom-built water baths [19,20,26], and outperforms PCB- and thin-film heaters, which often show deviations greater than ±1 °C and require 5–10 minutes to reach equilibrium [17,28,29,31]. The trade-off is that the water bath requires 10–15 minutes to reach the set temperature (65 °C), and cartridges containing µPAD strips or holder containing tubes can only be placed once set temperature is reached. In addition, the system requires water replacement periodically on weekly basis, whereas PCB heaters are essentially maintenance-free.

For illumination and color sensing, AquaStream uses backlit illumination with optimized camera settings for both µPADs (gain 2, 100 ms exposure) and tubes (gain 2, 10 ms exposure) (Figs 3B–D). This design choice stemmed from our prior experience with FARM-LAMP [19], where front illumination often produced shadows and reflection artifacts when imaging through a water bath. We evaluated the effect of light intensity by adjusting the camera gain, and under these conditions the HSV hue channel remained a reliable metric for tracking color transitions, with variability between replicates generally within ±2 hue units and a net hue difference of up to 50 between negative and positive reactions (pink to yellow transition, Figs 3B-D). However, hue values shifted systematically with gain settings, indicating that in backlit imaging, hue varies with lighting, unlike under front illumination [17,43]. This discrepancy likely reflects the fact that absorbance dominates in backlit imaging, whereas reflectance dominates in front illumination, leading to change in the hue values. Thus, while hue remains a robust choice for within-run analysis, careful control of gain and camera parameters is required for reproducibility across runs and instruments.

AquaStream achieves an effective balance between portability, throughput, and cost. The research prototype can be fabricated with an estimated BOM component cost of approximately USD 327 using 3D-printed parts and consumer-grade electronics. This estimated component cost is comparable to other reported low-cost analyzers such as FARM-LAMP [19], UbiNAAT [29], MILP [22] which are reported in the USD 200–1500 range, although the final commercial sale price of such instruments may be higher.. Despite its affordability, AquaStream supports up to 24 tube reactions or 42 µPAD zones per run, three to five times higher throughput than comparable systems. This scalability distinguishes it from most portable platforms, which typically accommodate fewer than eight tubes or a dozen µPADs. The only higher-throughput alternative is our previous ThermiQuant™ MegaScan, capable of 160 µPADs but restricted to paper assays, twice heavier, and nearly five times more costly [20]. The main trade-off is its 3 L water bath, which increases weight (∼5 kg) and reduces portability compared with ultra-miniaturized PCB-integrated systems such as [18]. Nonetheless, AquaStream’s combination of high throughput, low cost, and flexible format compatibility makes it particularly suited for decentralized testing and field deployment.

For onboard analysis, AquaStream integrates a software pipeline directly on the device, automating the full imaging workflow. It accurately detects µPAD zones and tubes (100% for µPADs and 99.7% for tubes at a 50% confidence threshold; Figs 2B–C), filters and averages hue values to minimize background effects, and generates hue–time trajectories that closely resemble quantitative PCR (qPCR)-style kinetics: sigmoidal curves for positives and linear traces for negatives (Figs 4–5) [44]. The few misclassifications observed in liquid assays (0.3%) occurred when large droplets split and adhered to the upper tube wall, creating atypical color regions. This rare failure mode would also challenge conventional threshold-based methods, which in such cases would likely register two separate objects or misidentify background artifacts as signal. Compared to prior systems that rely on smartphone apps for imaging and processing [17,29], or systems that require exporting images for external processing, including our own Amplimetrics™ pipeline [14], AquaStream™ performs all processing onboard. This approach minimizes hands-on effort, enables streamlined real-time analysis, and eliminates the need for post-run image transfer. In addition, the integrated Raspberry Pi features an onboard antenna with WiFi connectivity, allowing the device to be remotely accessed. This capability not only facilitates convenient data transfer but is also beneficial for remote monitoring, troubleshooting, and debugging.

Taken together, AquaStream enabled direct comparison of LAMP reactions in µPADs and tubes under identical experimental conditions and within a single workflow (Fig 6). Liquid reactions amplified ∼8.7 minutes faster than paper under identical conditions. We hypothesize that this delay may be attributable to the slower reagent transport and diffusion constraints inherent to cellulose matrices [45–47]. What AquaStream uniquely allowed, however, was a quantitative assessment of this difference, something that has not been possible with fragmented workflows.

The LOQ was defined as the lowest concentration at or above the LOD with CV ≤ 10%, provided that the next two consecutive higher concentrations also had CV ≤ 10%. CV was estimated only for concentrations with at least three numeric replicates. Therefore, LOQ determination required three consecutive concentrations meeting the CV threshold, starting from the candidate LOQ concentration. A 10% CV threshold was selected based on prior reports indicating that LAMP assays typically show acceptable repeatability within a CV range of approximately 5–10% [37–40]. This approach however has limitations. The choice of a CV threshold remains somewhat arbitrary because there is no universally accepted consensus for defining acceptable quantitative precision in LAMP assays; most reported values represent recommendations or best-practice guidelines rather than formal standards [36,48]. In the current study, we do not aim to develop a diagnostic but to demonstrate that AquaStream can characterize assay performance adequately. Other studies could use their own CV thresholds for LOQ depending on the technical capabilities of the assay and the clinical relevance of the diagnostic target. Another commonly held assumption is that analytical variability increases as analyte concentration decreases, leading to higher CV values at lower concentrations [36,48]. However, our data did not consistently follow this pattern. As shown in S6 Fig, CV values at higher concentrations (1E6 and 1E5 copies/reaction) were greater than those observed for several lower concentrations. In addition, isolated concentrations within the middle of the standard curve occasionally exhibited elevated CV values (S7 Fig), even when surrounding concentrations showed acceptable variability. Such local fluctuations complicate LOQ determination if it is based solely on identifying a single concentration meeting the CV threshold. We therefore applied a rule whereby the candidate LOQ concentration and the next two consecutive higher concentrations must all satisfy the CV ≤ 10% criterion for the lowest concentration to be considered the LOQ.

Overall, AquaStream is useful for applications that require relative comparison, whether within the same format or across formats. Although not demonstrated in this study, the platform could be leveraged for many future investigations. For example, there is growing interest in lyophilizing or drying reagents directly on µPADs or tubes, and AquaStream could be used to systematically evaluate how ageing or environmental conditions affect reaction kinetics over time [3,5,6,49]. Similarly, the effect of complex sample matrices such as blood or saliva on LAMP performance remains an area of active research [50–52]. AquaStream would enable direct comparison of these matrix effects against purified DNA targets, as well as reveal how such effects differ between liquid and paper-based assays. In this way, the platform provides researchers with a practical tool to explore reagent stability, matrix interference, and assay robustness under conditions that are directly relevant to real-world deployment.

## Conclusions

In this study, we introduced ThermiQuant™ AquaStream, a portable, low-cost water-bath incubator with integrated imaging and automated analysis for dual-format colorimetric NAATs on both µPADs and tubes. The system offers five major advantages: (i) dual-modality support, enabling both µPAD- and tube-based reactions within a single instrument, (ii) fully onboard software with automatic object detection and hue-trajectory analysis, reducing reliance on external devices and simplifying operation for users, (iii) precise and uniform isothermal heating at 65 ± 0.5 °C, (iv) higher throughput at lower cost than most comparable systems while retaining a compact benchtop footprint, and (v) an optimized backlit illumination and analysis strategy that enables reproducible hue-based kinetics and direct benchmarking between liquid and paper assays within a single workflow.

The platform also has two limitations: (i) the 3 L water bath increases overall weight (∼5 kg) and requires weekly replacement, making AquaStream bulkier and less portable than ultra-compact PCB-based systems, and (ii) the backlit illumination strategy, while improving image quality, makes hue measurements sensitive to camera gain and exposure, requiring strict parameter standardization to ensure reproducibility across runs and instruments.

Future work will address these gaps by using the following three approaches: (i) replacing the water bath with transparent heaters such as indium tin oxide (ITO)-coated glass heaters, optimized to achieve the ±1 °C precision required for reproducible LAMP while reducing weight and maintenance, (ii) developing mathematical and color correction models to create backlight-intensity-agnostic color metrics that minimize sensitivity to camera settings, and (iii) extending testing beyond synthetic DNA to include diverse biological samples such as blood, saliva, and environmental matrices, enabling robust performance benchmarking in realistic diagnostic scenarios.

## Supporting information

**S1 File. Supporting information document.** All supporting figures and tables are included.

**S2 File. Design Files.** All CAD design files are included.

**S3 File. Software Files.** Amplimetrics™ software as well as ThermiQuant™ AquaStream source codes.

## Data Availability

Raw and processed data presented here are available on Mendeley Data (DOI: 10.17632/2fhysyyrdc.2).

## Acknowledgements

We are grateful to Dr. Bilal Ahmed, Dr. Simerdeep Kaur, Ashley Kayabasi, Dr. Mohamed Kamel, Dr. Rachel Munds, Cindy Mayorga, Nafisa Rafiq, Dr. Dan Van Nguyen, for their participation in the initial user testing of the hardware and software tools and for providing insightful feedback. In addition, we are grateful for feedback on design and usability of the devices built here to the following people: Aaron Ault, Aaron Gilbertie, Dr. J. Alex Pasternak, Dr. Jon P Schoonmaker, Dr. Patrick Zollner, Dr. Arezoo Ardekani, Dr. Darryl Ragland, and Dr. Timothy Johnson.

## Author Contributions

**Bibek Raut**: Conceptualization, software, data curation, formal analysis, investigation, methodology, validation, visualization, writing – original draft preparation. **Gopal Palla**: Data curation, formal analysis, investigation, methodology, validation. **Jiangshan Wang**: Conceptualization, validation, and writing - review & editing. **Skyler Campbell**: Software, investigation, methodology. **Virendra Kumar**: Methodology, writing – review and editing. **Kyungyeon Ra**: Methodology, writing – review and editing. **Mohit S. Verma**: Conceptualization, supervision, project administration, funding acquisition, validation, and writing – review & editing. All authors have reviewed the manuscript and approved the final version.

## Competing Interest

Mohit S. Verma has the following interests in Krishi, Inc.: ownership, board membership, and research grants. Mohit S. Verma has the following interests in Simply Experiment LLC: ownership. Krishi, Inc. and Simply Experiment LLC did not fund this work. The authors would also like to declare the following patent application associated with this research: 19/010,865. Mohit S. Verma and Bibek Raut are inventors on the patent, which is assigned to Purdue Research Foundation.

## Use of Generative AI

During the preparation of this work, the authors used Grammarly and ChatGPT to check for grammar errors and improve their academic writing language as well as debugging the software. After using this tool/service, the authors reviewed and edited the content as needed and take full responsibility for the content of the publication.

## Copyright claim

ThermiQuant™ and Amplimetrics™ are copyrighted by © Purdue Research Foundation 2024.

## Notes

https://data.mendeley.com/datasets/2fhysyyrdc/1

### Summary of Updates

The manuscript has additional data to make the conclusions stronger and clearer.

### Funding

- Center For Produce Safety: 2023CPS11
- Agricultural Marketing Service: AM22SCBPCA1133
- Animal and Plant Health Inspection Service: AP22VSD&B000C022, AP23OA000000C015
- Purdue University West Lafayette: 2025 Bridge Funding

## References

1. Notomi T, Okayama H, Masubuchi H, Yonekawa T, Watanabe K, Amino N, et al. Loop-mediated isothermal amplification of DNA. Nucleic Acids Research. 2000;28: e63. doi:10.1093/nar/28.12.e63

2. Durward E, Harris WJ. Colorimetric Method for Detecting Amplified Nucleic Acids. BioTechniques. 1998;25: 608–614. doi:10.2144/98254st01

3. Sritong N, Medeiros MS de, Anthony Basing L, C. Linnes J. Promise and perils of paper-based point-of-care nucleic acid detection for endemic and pandemic pathogens. Lab on a Chip. 2023;23: 888–912. doi:10.1039/D2LC00554A

4. Thekkudan Novi V, Kumar Meher A, Abbas A. Visualization methods for loop mediated isothermal amplification (LAMP) assays. Analyst. 2025;150: 588–599. doi:10.1039/D4AN01287A

5. Das D, Masetty M, Priye A. Paper-Based Loop Mediated Isothermal Amplification (LAMP) Platforms: Integrating the Versatility of Paper Microfluidics with Accuracy of Nucleic Acid Amplification Tests. Chemosensors. 2023;11: 163. doi:10.3390/chemosensors11030163

6. Wang J, Davidson JL, Kaur S, Dextre AA, Ranjbaran M, Kamel MS, et al. Paper-Based Biosensors for the Detection of Nucleic Acids from Pathogens. Biosensors (Basel). 2022;12: 1094. doi:10.3390/bios12121094

7. Davidson JL, Wang J, Maruthamuthu MK, Dextre A, Pascual-Garrigos A, Mohan S, et al. A paper-based colorimetric molecular test for SARS-CoV-2 in saliva. Biosensors and Bioelectronics: X. 2021;9: 100076. doi:10.1016/j.biosx.2021.100076

8. Kamel M, Davidson JL, Schober JM, Fraley GS, Verma MS. A paper-based loop-mediated isothermal amplification assay for highly pathogenic avian influenza. Sci Rep. 2025;15: 12110. doi:10.1038/s41598-025-95452-6

9. Mori Y, Notomi T. Loop-mediated isothermal amplification (LAMP): a rapid, accurate, and cost-effective diagnostic method for infectious diseases. J Infect Chemother. 2009;15: 62–69. doi:10.1007/s10156-009-0669-9

10. Tanner NA, Zhang Y, Evans Jr. TC. Visual Detection of Isothermal Nucleic Acid Amplification Using pH-Sensitive Dyes. BioTechniques. 2015;58: 59–68. doi:10.2144/000114253

11. Zhang M, Liu J, Shen Z, Liu Y, Song Y, Liang Y, et al. A newly developed paper embedded microchip based on LAMP for rapid multiple detections of foodborne pathogens. BMC Microbiol. 2021;21: 197. doi:10.1186/s12866-021-02223-0

12. Aoki MN, de Oliveira Coelho B, Góes LGB, Minoprio P, Durigon EL, Morello LG, et al. Colorimetric RT-LAMP SARS-CoV-2 diagnostic sensitivity relies on color interpretation and viral load. Sci Rep. 2021;11: 9026. doi:10.1038/s41598-021-88506-y

13. Wong NCK, Meshkinfamfard S, Turbé V, Whitaker M, Moshe M, Bardanzellu A, et al. Machine learning to support visual auditing of home-based lateral flow immunoassay self-test results for SARS-CoV-2 antibodies. Commun Med. 2022;2: 78. doi:10.1038/s43856-022-00146-z

14. Ahmed B, Raut B, Pauley A, Davidson JL, Yang S, Verma MS. Development of a portable paper-based biosensor for the identification of genetically modified corn (*Zea mays*) and soybean (*Glycine max*). Biosensors and Bioelectronics. 2025;287: 117690. doi:10.1016/j.bios.2025.117690

15. Kaur M, Ayarnah K, Duanis-Assaf D, Alkan N, Eltzov E. Paper-based colorimetric loop-mediated isothermal amplification (LAMP) assay for the identification of latent *Colletotrichum* in harvested fruit. Analytica Chimica Acta. 2023;1267: 341394. doi:10.1016/j.aca.2023.341394

16. Seok Y, Joung H-A, Byun J-Y, Jeon H-S, Shin SJ, Kim S, et al. A Paper-Based Device for Performing Loop-Mediated Isothermal Amplification with Real-Time Simultaneous Detection of Multiple DNA Targets. Theranostics. 2017;7: 2220–2230. doi:10.7150/thno.18675

17. Nguyen HQ, Nguyen VD, Van Nguyen H, Seo TS. Quantification of colorimetric isothermal amplification on the smartphone and its open-source app for point-of-care pathogen detection. Sci Rep. 2020;10: 15123. doi:10.1038/s41598-020-72095-3

18. Papadakis G, Pantazis AK, Fikas N, Chatziioannidou S, Tsiakalou V, Michaelidou K, et al. Portable real-time colorimetric LAMP-device for rapid quantitative detection of nucleic acids in crude samples. Sci Rep. 2022;12: 3775. doi:10.1038/s41598-022-06632-7

19. Wang J, Kaur S, Kayabasi A, Ranjbaran M, Rath I, Benschikovski I, et al. A portable, easy-to-use paper-based biosensor for rapid in-field detection of fecal contamination on fresh produce farms. Biosensors and Bioelectronics. 2024;259: 116374. doi:10.1016/j.bios.2024.116374

20. Raut B, Palla G, Kumar V, Fleck A, Ahmed B, Davidson JL, et al. ThermiQuant(TM) MegaScan: High-throughput isothermal reactor with quantitative colorimetric readout for paper-based nucleic acid amplification tests. bioRxiv; 2026. p. 2026.01.09.696240. doi:10.64898/2026.01.09.696240

21. Jiang KP, Bennett S, Heiniger EK, Kumar S, Yager P. UbiNAAT: a multiplexed point-of-care nucleic acid diagnostic platform for rapid at-home pathogen detection. Lab Chip. 2024;24: 492–504. doi:10.1039/D3LC00753G

22. Kumar N, Kumari M, Chander D, Dogra S, Chaubey A, Chakraborty S, et al. Portable, quantitative, real-time isothermal nucleic acid amplification test using microfluidic device-coupled UV-LED photodiode detector. Biosensors and Bioelectronics. 2025;274: 117194. doi:10.1016/j.bios.2025.117194

23. Cui S, Wang K, Yang Y, Lv X, Li X. An integrated and paper-based microfluidic system employing LAMP-CRISPR and equipped with a portable device for simultaneous detection of pathogens. Anal Bioanal Chem. 2025;417: 785–797. doi:10.1007/s00216-024-05693-z

24. Bai H, Liu Y, Gao L, Wang T, Zhang X, Hu J, et al. A portable all-in-one microfluidic device with real-time colorimetric LAMP for HPV16 and HPV18 DNA point-of-care testing. Biosensors and Bioelectronics. 2024;248: 115968. doi:10.1016/j.bios.2023.115968

25. Pascual-Garrigos A, Maruthamuthu MK, Ault A, Davidson JL, Rudakov G, Pillai D, et al. On-farm colorimetric detection of Pasteurella multocida, Mannheimia haemolytica, and Histophilus somni in crude bovine nasal samples. Vet Res. 2021;52: 126. doi:10.1186/s13567-021-00997-9

26. Rafiq N, Verma MS. Design and Development of a Field-Deployable Water Bath for Loop-Mediated Isothermal Amplification Assay. IEEE Sensors Journal. 2025; 1–1. doi:10.1109/JSEN.2025.3588790

27. Myers FB, Moffatt B, El Khaja R, Chatterjee T, Marwaha G, McGee M, et al. A robust, low-cost instrument for real-time colorimetric isothermal nucleic acid amplification. PLoS One. 2022;17: e0256789. doi:10.1371/journal.pone.0256789

28. García-Bernalt Diego J, Fernández-Soto P, Márquez-Sánchez S, Santos Santos D, Febrer-Sendra B, Crego-Vicente B, et al. SMART-LAMP: A Smartphone-Operated Handheld Device for Real-Time Colorimetric Point-of-Care Diagnosis of Infectious Diseases via Loop-Mediated Isothermal Amplification. Biosensors. 2022;12: 424. doi:10.3390/bios12060424

29. Jiang KP, Bennett S, Heiniger EK, Kumar S, Yager P. UbiNAAT: a multiplexed point-of-care nucleic acid diagnostic platform for rapid at-home pathogen detection. Lab Chip. 2024;24: 492–504. doi:10.1039/D3LC00753G

30. K. Heiniger E, P. Jiang K, Kumar S, Yager P. A low-cost point-of-care device for the simultaneous detection of two sexually transmitted bacterial pathogens in vaginal swab samples. Analyst. 2025;150: 4414–4426. doi:10.1039/D5AN00496A

31. Cui S, Wang K, Yang Y, Lv X, Li X. An integrated and paper-based microfluidic system employing LAMP-CRISPR and equipped with a portable device for simultaneous detection of pathogens. Anal Bioanal Chem. 2025;417: 785–797. doi:10.1007/s00216-024-05693-z

32. Wang J, Dextre A, Pascual-Garrigos A, Davidson JL, Maruthamuthu MK, McChesney D, et al. Fabrication of a paper-based colorimetric molecular test for SARS-CoV-2. MethodsX. 2021;8: 101586. doi:10.1016/j.mex.2021.101586

33. Varghese R, M. S. YOLOv8: A Novel Object Detection Algorithm with Enhanced Performance and Robustness. 2024 International Conference on Advances in Data Engineering and Intelligent Computing Systems (ADICS). 2024. pp. 1–6. doi:10.1109/ADICS58448.2024.10533619

34. Stokdyk JP, Firnstahl AD, Spencer SK, Burch TR, Borchardt MA. Determining the 95% limit of detection for waterborne pathogen analyses from primary concentration to qPCR. Water Research. 2016;96: 105–113. doi:10.1016/j.watres.2016.03.026

35. Blommel JH, Jenkinson G, Binnicker MJ, Karon BS, Boccuto L, Ivankovic DS, et al. Authorized SARS-CoV-2 molecular methods show wide variability in the limit of detection. Diagnostic Microbiology and Infectious Disease. 2023;105: 115880. doi:10.1016/j.diagmicrobio.2022.115880

36. Klymus KE, Merkes CM, Allison MJ, Goldberg CS, Helbing CC, Hunter ME, et al. Reporting the limits of detection and quantification for environmental DNA assays. Environmental DNA. 2020;2: 271–282. doi:10.1002/edn3.29

37. Lou Y, Chen Y, Zhou Q. Simultaneous detection of multiple spoilage bacteria using a fluorogenic loop-mediated isothermal amplification-based multi-sample microfluidic chip. LWT. 2024;193: 115758. doi:10.1016/j.lwt.2024.115758

38. Hayes EK, Gouthro MT, Gagnon GA. Isothermal amplification as a water safety tool: rapid detection of viruses in surface water and wastewater. Environ Sci: Water Res Technol. 2025;11: 2141–2151. doi:10.1039/D5EW00092K

39. Lalonde L, Xie V, Lobanov V. Optimization and validation of a loop-mediated isothermal amplification (LAMP) assay for detection of Giardia duodenalis in leafy greens. Food and Waterborne Parasitology. 2021;23: e00123. doi:10.1016/j.fawpar.2021.e00123

40. Wu Y, Bai L, Ye C, Yuhong Guan, Kunming Yan, Chen H, et al. Novel miniaturized fluorescence loop-mediated isothermal amplification detection system for rapid on-site virus detection. Front Bioeng Biotechnol. 2022;10. doi:10.3389/fbioe.2022.964244

41. Garcia-Torales G, Torres-Ortega HH, Estrada-Marmolejo R, Beltran-Gonzalez AB, Strojnik M. Thermal Bed Design for Temperature-Controlled DNA Amplification Using Optoelectronic Sensors. Sensors. 2024;24: 7050. doi:10.3390/s24217050

42. Matusiak T, Tokarski M, Małodobra-Mazur M, Roguszczak H, Dąbrowski A, Sitarz P, et al. Contactless Heating Technology for Lab-on-Chip Microfluidic-Based Nucleic Acid Amplification Testing System. Proceedings. 2024;97: 58. doi:10.3390/proceedings2024097058

43. Yin K, Pandian V, Kadimisetty K, Zhang X, Ruiz C, Cooper K, et al. Real-time Colorimetric Quantitative Molecular Detection of Infectious Diseases on Smartphone-based Diagnostic Platform. Sci Rep. 2020;10:9009. doi:10.1038/s41598-020-65899-w

44. Zhang Y, Li H, Shang S, Meng S, Lin T, Zhang Y, et al. Evaluation validation of a qPCR curve analysis method and conventional approaches. BMC Genomics. 2021;22: 680. doi:10.1186/s12864-021-07986-4

45. Chaudhury K, Kar S, Chakraborty S. Diffusive dynamics on paper matrix. Appl Phys Lett. 2016;109: 224101. doi:10.1063/1.4966992

46. Das D, Panigrahi PK. CFD simulations for paper-based DNA amplification reaction (LAMP) of Mycobacterium tuberculosis—point-of-care diagnostic perspective. Med Biol Eng Comput. 2020;58: 271–289. doi:10.1007/s11517-019-02082-y

47. Elizalde E, Urteaga R, Berli CLA. Rational design of capillary-driven flows for paper-based microfluidics. Lab Chip. 2015;15: 2173–2180. doi:10.1039/C4LC01487A

48. Forootan A, Sjöback R, Björkman J, Sjögreen B, Linz L, Kubista M. Methods to determine limit of detection and limit of quantification in quantitative real-time PCR (qPCR). Biomol Detect Quantif. 2017;12: 1–6. doi:10.1016/j.bdq.2017.04.001

49. Kang T, Lu J, Yu T, Long Y, Liu G. Advances in nucleic acid amplification techniques (NAATs): COVID-19 point-of-care diagnostics as an example. Biosensors and Bioelectronics. 2022;206: 114109. doi:10.1016/j.bios.2022.114109

50. Moehling TJ, Choi,Gihoon, Dugan,Lawrence C., Salit,Marc, and Meagher RJ. LAMP Diagnostics at the Point-of-Care: Emerging Trends and Perspectives for the Developer Community. Expert Review of Molecular Diagnostics. 2021;21: 43–61. doi:10.1080/14737159.2021.1873769

51. Nwe MK, Jangpromma N, Taemaitree L. Evaluation of molecular inhibitors of loop-mediated isothermal amplification (LAMP). Sci Rep. 2024;14: 5916. doi:10.1038/s41598-024-55241-z

52. Selva Sharma A, Lee NY. Advancements in visualizing loop-mediated isothermal amplification (LAMP) reactions: A comprehensive review of colorimetric and fluorometric detection strategies for precise diagnosis of infectious diseases. Coordination Chemistry Reviews. 2024;509: 215769. doi:10.1016/j.ccr.2024.215769

