## Supplementary File 1 for "ThermiQuant™ AquaStream: A portable instrument for quantitative colorimetric isothermal nucleic acid amplification reactions in paper and tube formats"

for

*Corresponding author

### 1. Supporting Figures

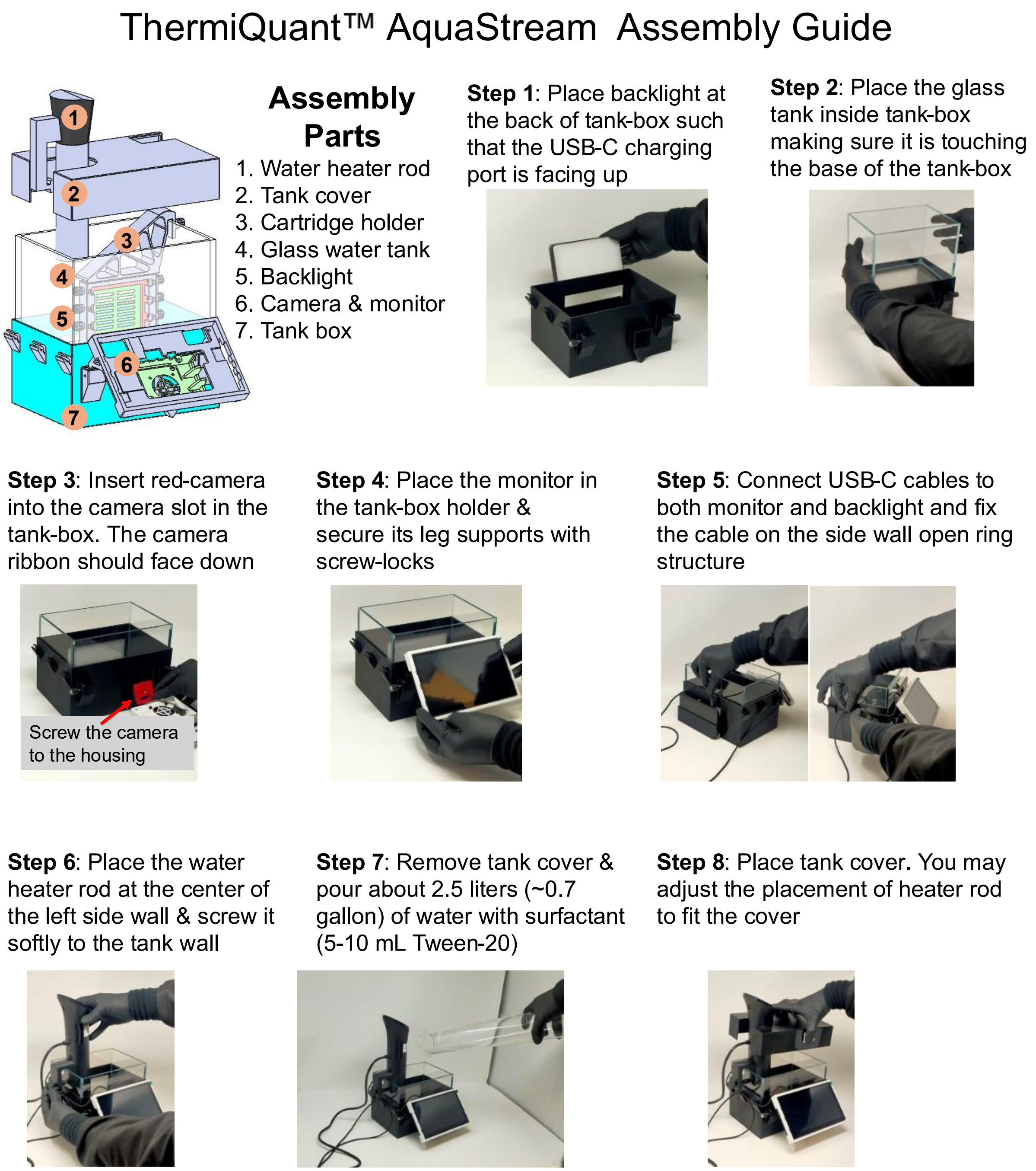

**S1 Fig. Step-by-step visual guide to assemble ThermiQuant™ AquaStream for the first time.**

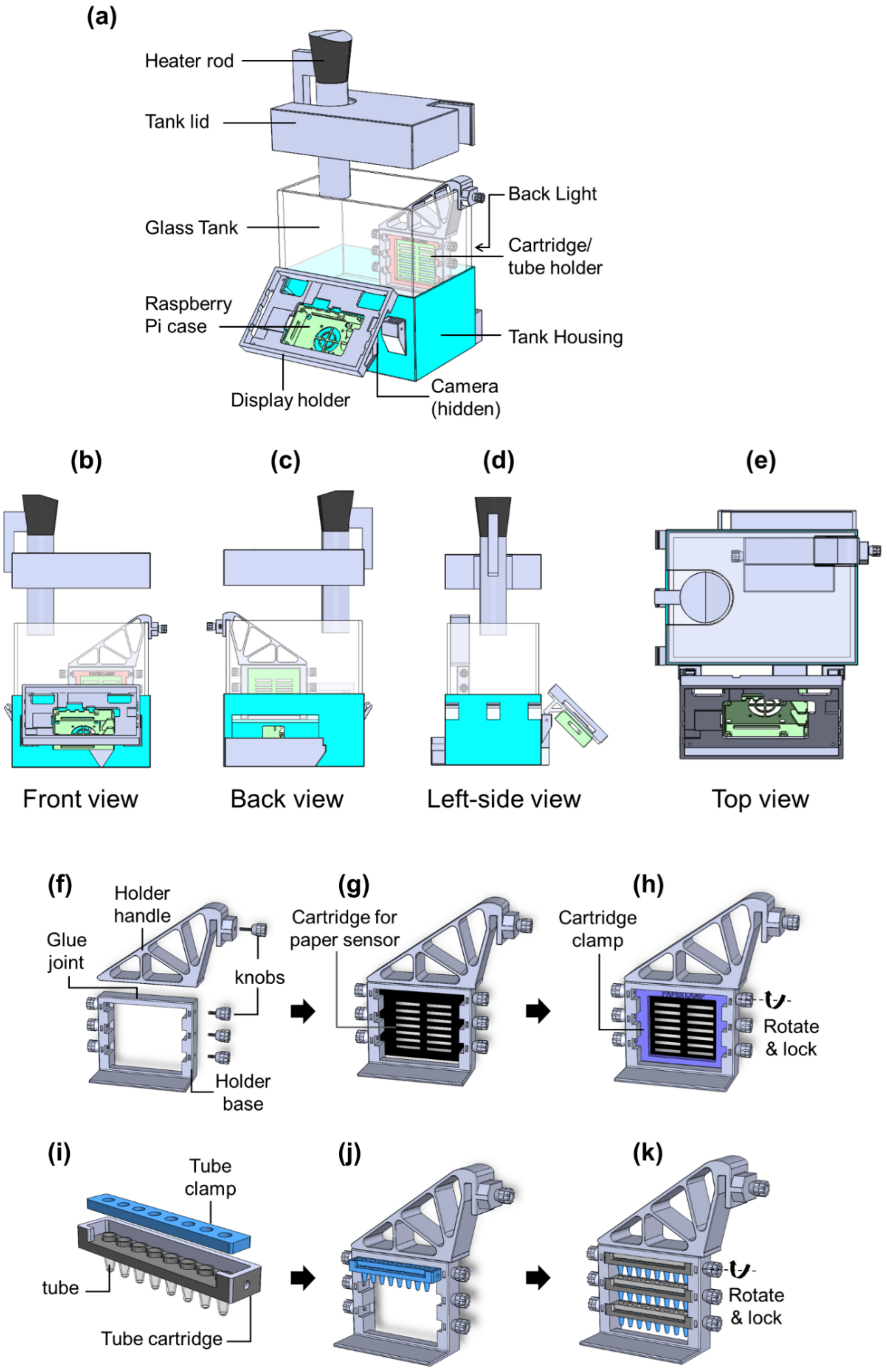
(a)
Heater rod
Tank lid
Glass Tank
Raspberry
Pi case
Display holder
Back Light
Cartridge/
tube holder
Tank Housing
Camera
(hidden)
(b)
(c)
(d)
(e)
Front view
Back view
Left-side view
Top view
(f)
Holder
handle
Glue
joint
knobs
Holder
base
(g)
Cartridge for
paper sensor
(h)
Cartridge
clamp
Rotate
& lock
(i)
Tube
clamp
tube
Tube cartridge
(j)
(k)
Rotate
& lock

**S2 Fig. 3D view of device and cartridge assembly.** (a) Exploded view of the instrument. (b) Front view. (c) Back view. (d) Left-side view. (e) Top view. (f) Exploded assembly view of the cartridge/tube holder showing two parts, the holder handle and the holder base, which need to be glued together using superglue. The knobs are fitted with M3 screws and nuts to secure the holder to the tank wall or to lock the cartridges in place. (g) To load the paper sensor cartridge, first place the cartridge, then (h) position the cartridge clamp and lock it with the knobs by rotating clockwise for the right side and counterclockwise for the left side. (i) Assembled view of the tube-inside-tube cartridge. (j) Insert the assembled tube cartridge into the cartridge holder, and (k) lock it in place before placing it inside the water bath.

### ThermiQuant™ AquaStream User Manual

**Step 1**: If not filled already, remove tank cover & pour about 2.5 liters (~0.7 gallon) of water with surfactant (10 mL Tween-20).

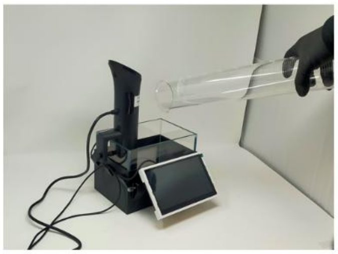

**Step 2**: Cover the tank & power the heater (set at 149 °F /65°C). Also, power the monitor & backlight. It takes ~10 min to heat water.

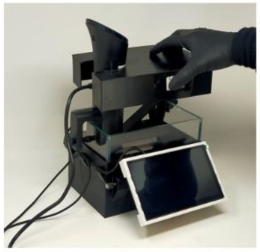

**Step 3A**: After water reaches 65°C, prepare to load the cartridge inside the holder by first inserting cartridge then adding a rectangular support.

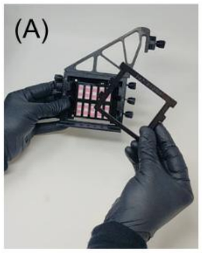

**Step 4B**: (B) Lock the cartridge & support using screw locks by rotating in opposite directions.

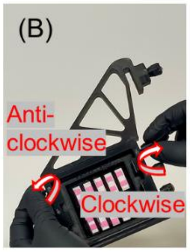

**Step 5A:** Gently insert the cartridge holder and make sure it is touching the back wall and lays flat on the tank.

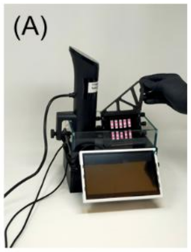

**Step 5B:** Then gently secure the cartridge to right side wall using screw lock. Do not tighten too much.

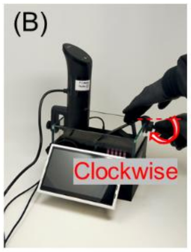

**Step 6.** Cover the tank with lid to eliminate outside light and reduce water evaporation.

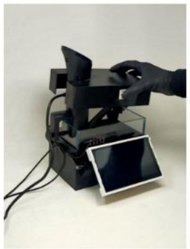

**Step 7.** In the software select the reaction type and press start to begin the reaction.

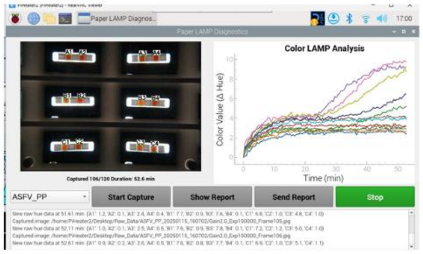

**Step 8.** After 60 minutes, a result summary screen appears that shows the reaction results with quantitative information.

| Label | Result | Tq (min) |
|---|---|---|
| A1 | Positive | 35 |
| A2 | Negative | -- |
| B1 | Positive | 30 |
| B3 | Negative | -- |

**S3 Fig. ThermiQuant™ AquaStream user manual.**

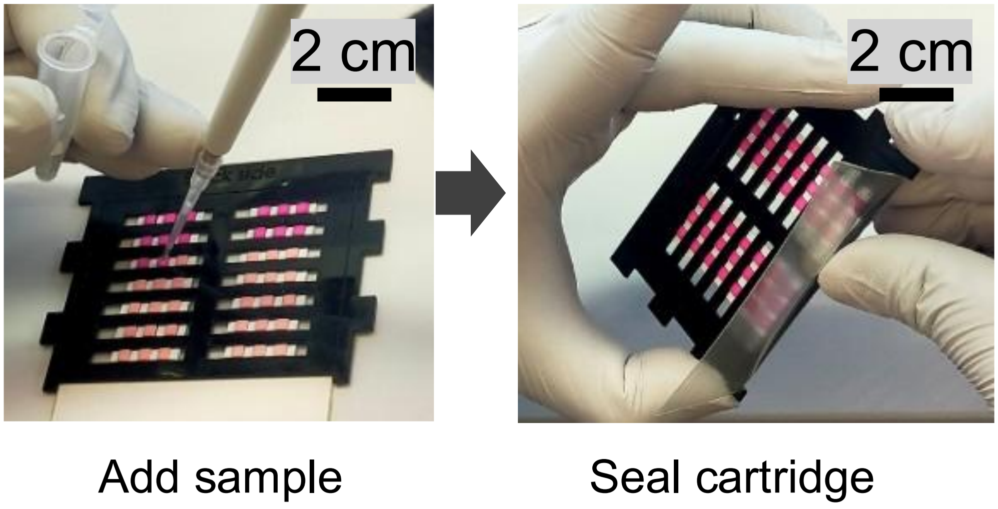

**S4 Fig. µPAD cartridge housing assembly.** Sample pipetting into µPADs pre-dried with reagents required for LAMP amplification, followed by sealing with PCR tape. Each µPAD zone was loaded with 7.5 µL of sample by pipette, after which the strip was immediately sealed with transparent PCR tape to prevent evaporation and cross-contamination during incubation.

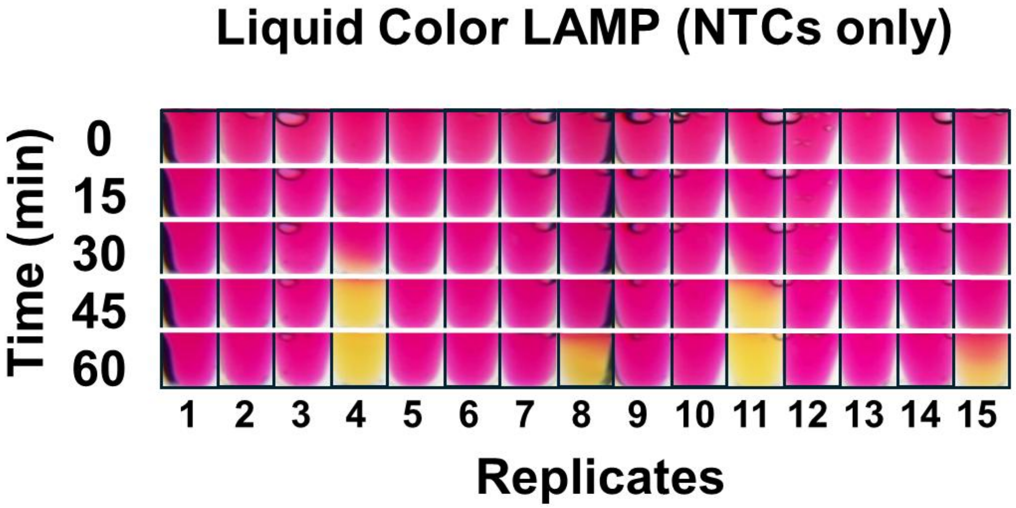

**S5 Fig. Timelapse of NTC-only reactions in tubes.** Liquid Color LAMP was used to test 15 additional NTC replicates to determine positivity threshold.

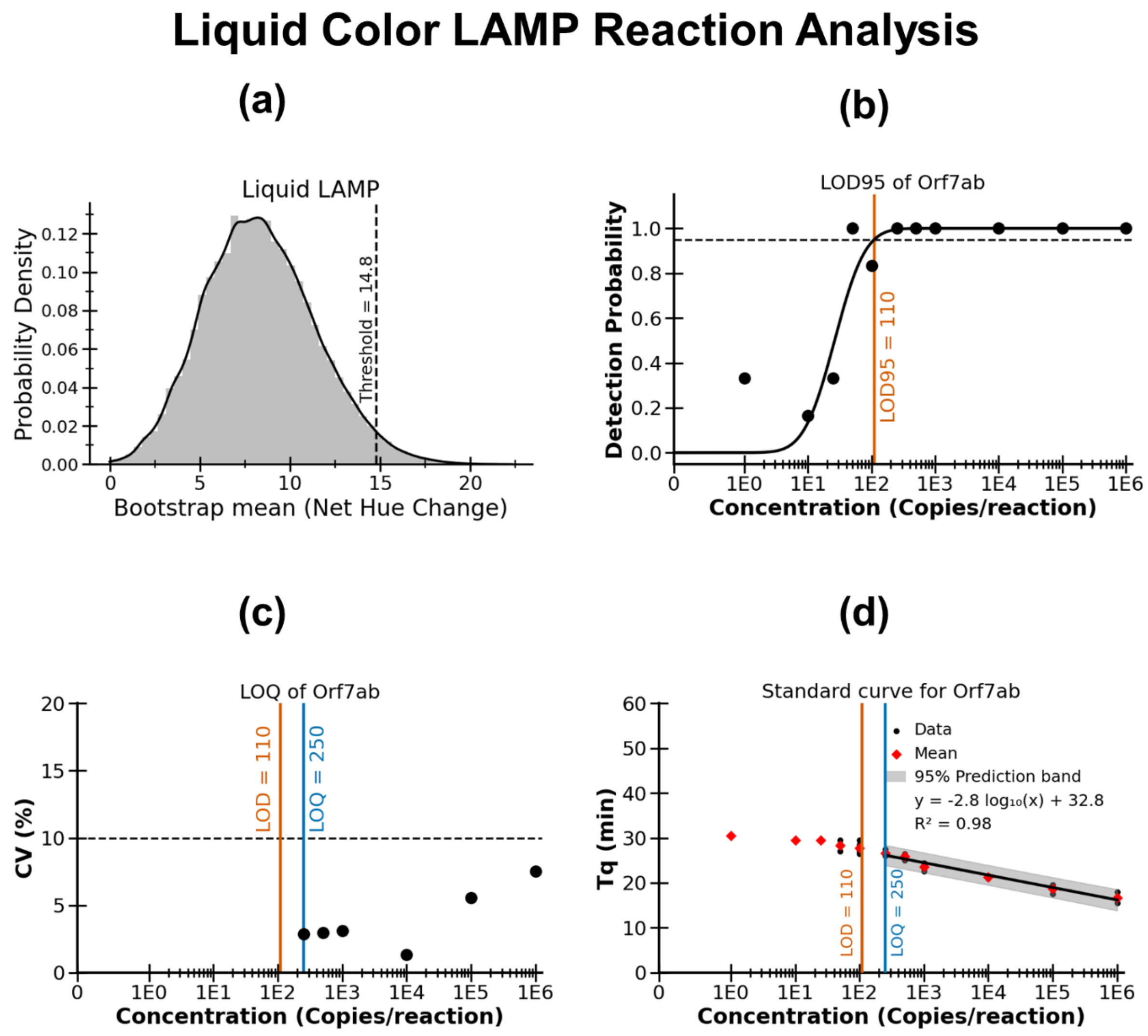

**S6 Fig. Determination of positivity threshold, LOD, and LOQ in liquid colorimetric LAMP.** (a) Probability density plot of the mean bootstrap net hue change derived from a total of n = 21 NTCs, used to determine the positivity threshold as the 95% upper confidence limit of the bootstrap mean. (b) Probit regression used to determine the LOD95 (95% detection probability). (c) LOQ defined as the lowest analytical concentration tested with a coefficient of variation (CV) ≤ 10% of the quantification time (Tq). (d) Standard curve generated using concentrations above the LOQ, also showing Tq mean and raw values for all positive samples, including those falling below the LOD95.

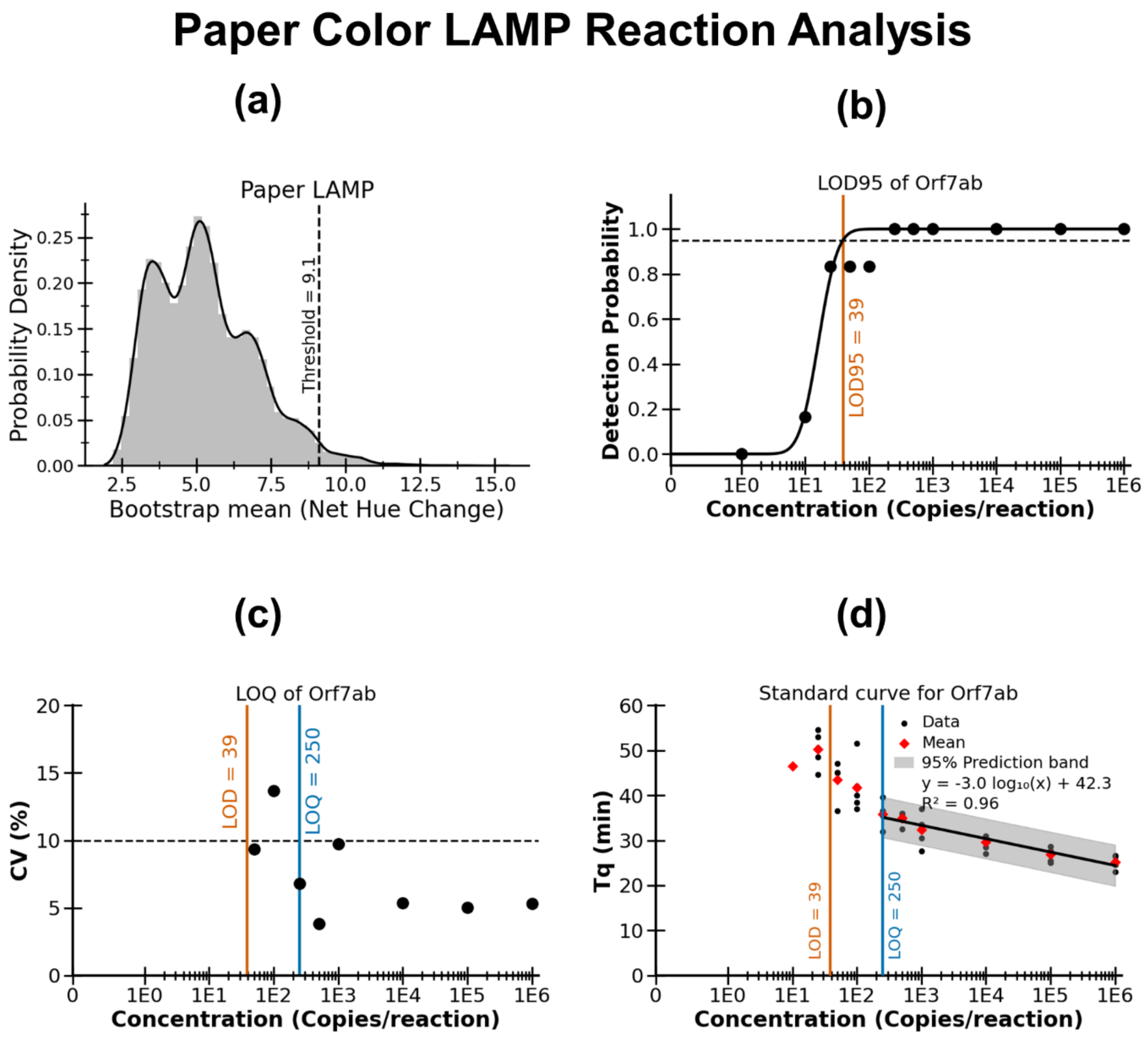

**S7 Fig. Determination of positivity threshold, LOD, and LOQ in paper colorimetric LAMP.** (a) Probability density plot of the mean bootstrap net hue change derived from a total of n = 14 NTCs, used to determine the positivity threshold as the 95% upper confidence limit of the bootstrap mean. (b) Probit regression used to determine the LOD95 (95% detection probability). (c) LOQ defined as the lowest analytical concentration tested with a coefficient of variation (CV) ≤ 10% of the quantification time (Tq). (d) Standard curve generated using concentrations above the LOQ, also showing Tq mean and raw values for all positive samples, including those falling below the LOD95.

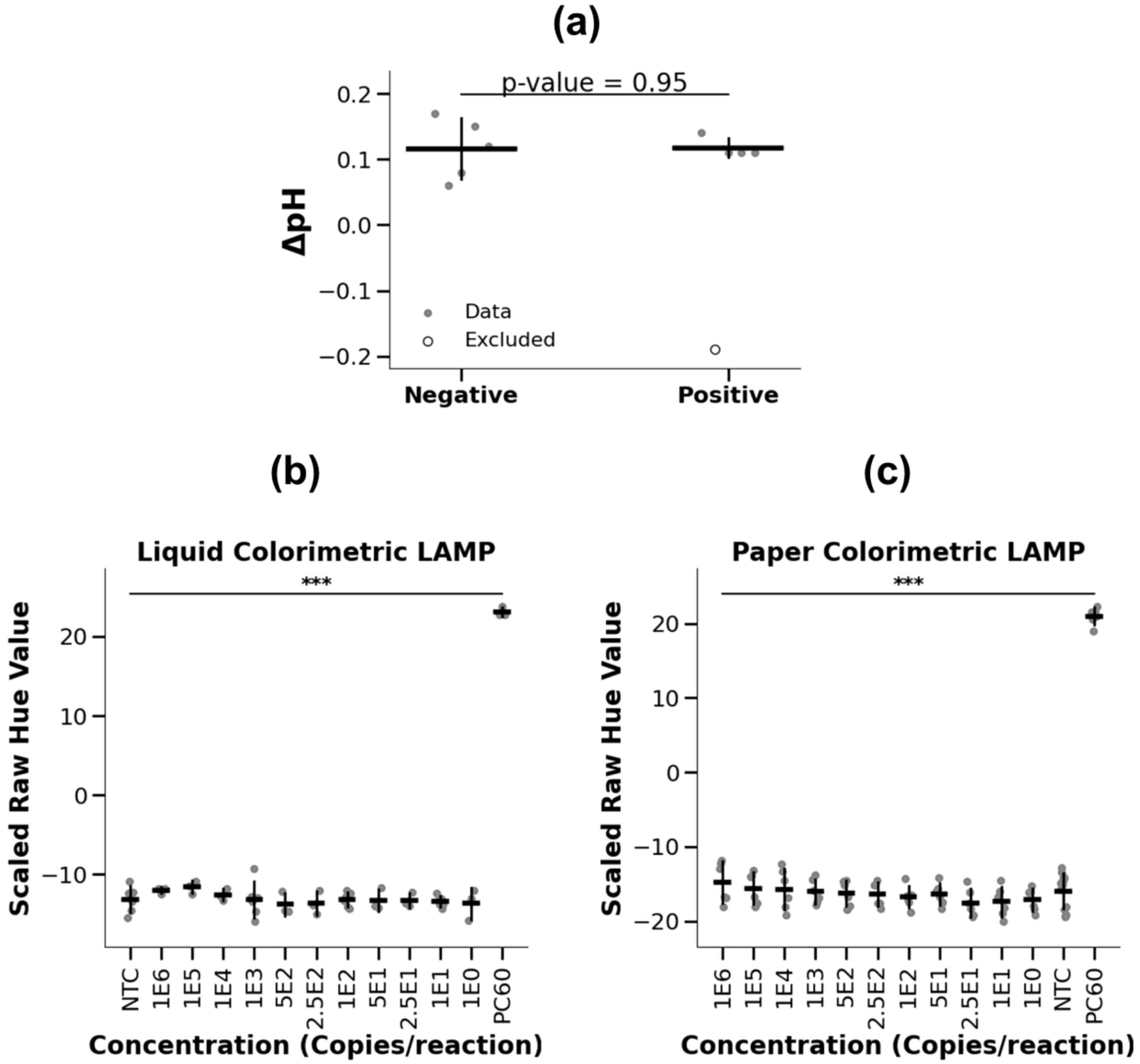

**S8 Fig. Effect of DNA concentration on the initial pH and hue value.** (A) Change in pH (ΔpH) between the first and second pH measurements for negative (water-only) and positive (DNA-spiked, 40,000 copies/µL) samples. Error bars represent mean ± SD. One positive replicate was excluded as indicated. No significant difference was observed (Welch's t-test, $p = 0.95$). (B) Raw hue values of liquid colorimetric LAMP reactions measured at the initial time point (0 min) across different DNA concentrations (copies per reaction). (C) Raw hue values of paper-based colorimetric LAMP reactions measured at 0 min across the same DNA concentrations. Error bars represent the mean ± SD. NTC denotes the no-template control measured at 0 min. PC60 denotes the positive control (1E6 copies per reaction) measured at 60 min after amplification. No significant differences in hue values were observed between the NTC at 0 min and any DNA concentration measured at 0 min. In contrast, a significant difference ($p < 0.001$) was observed between the NTC (0 min) and PC60.

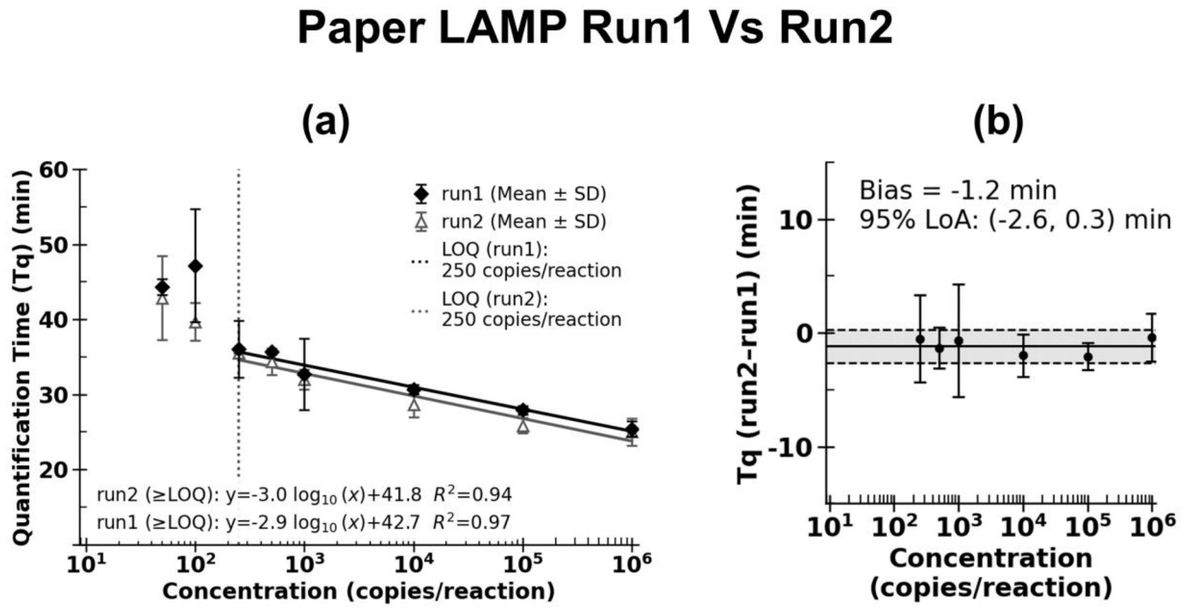

**S9 Fig. Inter-run variation in Tq of colorimetric paper LAMP assays.** (A) Quantification time (Tq) plotted against input DNA concentration (log scale) for µPAD assays in run 1 and run 2 at and above the LOD95, with linear regression performed only for concentrations at or above the LOQ. (B) Bland–Altman analysis comparing the differences in Tq between runs.

#### 2. Supporting Tables

**S1 Table. BOM of ThermiQuant™ MegaScan.** All reported prices are for Oct 2025.

**Bill of materials for the ThermiQuant™ AquaStream**

| S.N. | Parts Name | Manufacturer | Vendor | Vendor Parts Number | Units | Price/ Unit (USD) | Total Price (USD) |
|---|---|---|---|---|---|---|---|
| **1** | Raspberry Pi 4B (4GB) | Raspberry Pi Ltd | Digikey, USA | 2648-SC0194(9)-ND | 1 | 55 | 55 |
| **2** | microSD card (64 GB) | Amazon | Amazon, USA | ASIN: B08TJTB8XS | 1 | 8 | 7.54 |
| **3** | Power Supply Adapter (5V, 3A) | LitStar | Amazon, USA | ASIN: B08523QCT6 | 2 | 10 | 19.98 |
| **4** | Arducam 16 Megapixel camera | Arducam | Arducam, USA | IMX519 | 1 | 30 | 29.99 |
| **5** | Anova Nano 3.0 Sous Vide Precision Cooker | Anova | Amazon, USA | ASIN: B0BQ93XGWC | 1 | 72 | 72.35 |
| **6** | Small Nano Rimless Tank (1.1 gallon) | awxzom tank | Amazon, USA | ASIN: B0D3F36CTY | 1 | 29.99 | 29.99 |
| **7** | Touchscreen Monitor (7 inches, 800x480 pixel) | Freenove | Amazon, USA | ASIN: B0B44VZTRG | 1 | 49.95 | 49.95 |
| **8** | Rechargeable White LED Light (4000mAh) | Weilisi | Amazon, USA | ASIN: B09C5XKW64 | 1 | 29.99 | 29.99 |
| **9** | 3D Printer Filament (PETG and PC, 1 kg) | OVERTURE | Amazon, USA | ASIN: B0BQR8VGSR (PC White); B07PGYHYV8 (PETG Black) | 1 | 22.99 | 22.99 |
| **10** | M3 Hex socket head screw, spacer & nuts kit (stainless steel) | WZHUIDA | Amazon, USA | ASIN: B0CNJW37NJ | 1 | 8.99 | 8.99 |
| | | | | | | **Total** | **326.77** |

**Synthetic DNA, LAMP Primers, and qPCR probes and primers**

We used dPCR for absolute quantification, and LAMP was used for testing both µPAD and liquid-tube based assay. This study builds on a previously reported design of µPAD-based LAMP reactions targeting the *orf7ab* region of SARS-CoV-2[1–3].

**S2 Table. Orf7ab synthetic DNA sequence.**

| **Sequence (5'-3') target sequence of *orf7ab*** |
|---|
| ATGAAAATTATTCTTTTCTTGGCACTGATAACACTCGCTACTTGTGAGCTTTATCACT<br>ACCAAGAGTGTGTTAGAGGTACAACAGTACTTTTAAAAGAACCTT<br>GCTCTTCTGGAACATACGAGGGCAATTCACCATTTCATCCTCTAGCTGATAACAAAT<br>TTGCACTGACTTGCTTTAGCACTCAATTTGCTTTTGCTTGTCCTGAC<br>GGCGTAAAACACGTCTATCAGTTACGTGCCAGATCAGTTTCACCTAAACTGTTCATC<br>AGACAAGAGGAAGTTCAAGAACTTTACTCTCCAATTTTTCTTATTGT<br>TGCGGCAATAGTGTTTATAACACTTTGCTTCACACTCAAAAGAAAGACAGAATGAT<br>TGAACTTTCATTAATTGACTTCTATTTGTGCTTTTTAGCCTTTCTGCTATTCCTTGTTT<br>TAATTATGCTTATTATCTTTTGGTTCTCACTTGAACTGCAAGATCATAATGAAACTTG<br>TCACGCCTAA |

**S3 Table. Digital PCR (dPCR) primers and probes.** Same as previous report[3].

| **dPCR Primers** | **Sequence (5'-3')** | **Vendor** |
|---|---|---|
| orf7ab_PCR2_FWD | GAGGGCAATTCACCATTTCATC | Life Science Tech |
| orf7ab_PCR2_REV | AAACTGATCTGGCACGTAACT | Life Science Tech |
| Probe | /56-FAM/TT TGC TTG T/ZEN/C CTG ACG GCG TAA A/3IABkFQ/ | IDT |

**S4 Table. LAMP primers.** Same as previous report[3]

| **Primers** | **Sequence (5'-3')** | **Vendor** |
|---|---|---|
| SC2.orf7ab.1_F3 | CGGCGTAAAACACGTCTA | IDT |
| SC2.orf7ab.1_B3 | GCTAAAAAGCACAAATAGAAG TC | IDT |
| SC2.orf7ab.1_FIP | GGAGAGTAAAGTTCTTGAACTT CCTAGTTACGTGCCAGATCAG | IDT |
| SC2.orf7ab.1_BIP | TGCGGCAATAGTGTTTATAAC ACTATGAAAGTTCAATCATTCT GTCT | IDT |
| SC2.orf7ab.1_LF | TGTCTGATGAACAGTTTAGGT GAAA | IDT |
| SC2.orf7ab.1_LB | TTGCTTCACACTCAAAAGAA | IDT |

#### References

1. Davidson, J. L. *et al.* A paper-based colorimetric molecular test for SARS-CoV-2 in saliva. *Biosensors and Bioelectronics: X* **9**, 100076 (2021).
2. Wang, J. *et al.* Fabrication of a paper-based colorimetric molecular test for SARS-CoV-2. *MethodsX* **8**, 101586 (2021).
3. Raut, B. *et al.* ThermiQuant(TM) MegaScan: High-throughput isothermal reactor with quantitative colorimetric readout for paper-based nucleic acid amplification tests. 2026.01.09.696240 Preprint at https://doi.org/10.64898/2026.01.09.696240 (2026).
