## Supplementary File 3 for "ThermiQuant™ AquaStream: A portable instrument for quantitative colorimetric isothermal nucleic acid amplification reactions in paper and tube formats": LiquidLAMP.py

```
'''
Bibek Raut
Purdue University

Software to capture timelapse iamges. 
Settings are for Liquid LAMP. 
It captures images every 30 seconds for 60 minutes at camera gain 1,2,3,4,5. 
These settings can be customized. 

'''
from picamera2 import Picamera2, Preview
import time
import os
import datetime
from libcamera import controls
import math

# ---------------------- Global Constants ----------------------
CYCLES = 122               # Number of cycles
CYCLE_INTERVAL = 30        # Time for each cycle in seconds
SHORT_INTERVAL = 2         # Interval between each gain capture
GAIN_SETTINGS = [1.0,2.0,3.0,4.0,5.0]      # List of gains if you want multiple in one cycle
DISCARD_WAIT = 3           # Wait time before discarding first frame after start
INITIAL_SETUP_DURATION = 5 # Additional waiting for autofocus or other setups
EXPOSURE_TIME = 10000     # In microseconds, e.g., 10000 = 1/100s

def get_highest_numbered_folder(path):
    folder_numbers = []
    for folder_name in os.listdir(path):
        if os.path.isdir(os.path.join(path, folder_name)):
            try:
                number = int(folder_name.split('_')[-1])
                folder_numbers.append(number)
            except ValueError:
                pass
    return max(folder_numbers) if folder_numbers else 0

def make_new_folder():
    desktop_path = os.path.join(os.path.expanduser("~"), 'Desktop', 'Raw_Data')
    if not os.path.exists(desktop_path):
        os.makedirs(desktop_path, exist_ok=True)
        print("Created folder:", desktop_path)

    highest_number = get_highest_numbered_folder(desktop_path)
    current_date = datetime.datetime.now().strftime('%Y%m%d')
    folder_name = f"{current_date}_liqLAMP_{highest_number + 1}"
    folder_path = os.path.join(desktop_path, folder_name)
    
    os.makedirs(folder_path, exist_ok=True)
    print("Created folder:", folder_path)
    return folder_path

def main():
    # Create main folder
    main_folder_path = make_new_folder()

    # Create subfolders for each gain
    subfolders = {}
    frame_numbers = {}
    for gain in GAIN_SETTINGS:
        subfolder_name = f"Gain_{int(gain)}"
        subfolder_path = os.path.join(main_folder_path, subfolder_name)
        os.makedirs(subfolder_path, exist_ok=True)
        subfolders[gain] = subfolder_path
        frame_numbers[gain] = 1
        print("Created subfolder:", subfolder_path)

    # Initialize camera
    picam2 = Picamera2()
    initial_controls = {"AfMode": 1, "AfTrigger": 0}
    preview_config = picam2.create_preview_configuration(
        controls=initial_controls, main={"size": (1440, 1440)}, buffer_count=1
    )
    picam2.configure(preview_config)

    # Start camera preview
    picam2.start_preview(Preview.QTGL)
    picam2.start()
    print("Camera started.")

    # ----------------------------------------------------------
    # 1) Discard a frame immediately after camera starts
    # ----------------------------------------------------------
    print(f"Waiting {DISCARD_WAIT}s to let camera adjust, then discarding a frame...")
    time.sleep(DISCARD_WAIT)
    _ = picam2.capture_array()  # Captures to memory (not saved)
    print("Discarded the first camera frame (not saved).")

    # Extra time for initial setup or autofocus
    print(f"Waiting an additional {INITIAL_SETUP_DURATION}s for autofocus or stabilization...")
    time.sleep(INITIAL_SETUP_DURATION)

    # Optional: Trigger autofocus
    print("Triggering autofocus...")
    picam2.set_controls({"AfTrigger": 1})
    autofocus_attempts = 3
    lens_position = None
    for attempt in range(autofocus_attempts):
        print(f"Autofocus attempt {attempt+1}/{autofocus_attempts}")
        time.sleep(2)
        metadata = picam2.capture_metadata()
        lens_position = metadata.get("LensPosition")
        if lens_position is not None:
            print("Lens Position:", lens_position)
            break
    else:
        print("Autofocus did not find a lens position.")

    # Lock lens position if found
    if lens_position is not None:
        picam2.set_controls({"AfMode": 0, "LensPosition": lens_position})
        print("Lens position locked for manual focus.")

    print("Initial setup complete. Beginning timelapse cycles.\n")

    first_image_captured = False
    start_time = None

    # ----------------------------------------------------------
    # 2) Within each cycle, after setting exposure/gain,
    #    discard another "warm-up" frame before saving
    # ----------------------------------------------------------
    for cycle in range(CYCLES):
        cycle_start_time = time.time()
        print(f"Starting cycle {cycle+1}/{CYCLES} at {datetime.datetime.now()}")

        for gain in GAIN_SETTINGS:
            try:
                # Apply manual exposure/gain
                picam2.set_controls({"ExposureTime": EXPOSURE_TIME, "AnalogueGain": gain})
                time.sleep(0.3)  # Small wait to let the new settings apply

                # --- Discard a "warm-up" frame after new settings ---
                _ = picam2.capture_array()  # Not saved
                time.sleep(0.2)

                # --- Now capture the real image to file ---
                frame_number = frame_numbers[gain]
                file_name = f"Gain{int(gain)}_Exp{EXPOSURE_TIME}_Frame{frame_number}.jpg"
                file_path = os.path.join(subfolders[gain], file_name)
                picam2.capture_file(file_path)
                frame_numbers[gain] += 1

                print("Saved:", file_path)

                # Track overall elapsed time from the first official image
                if not first_image_captured:
                    start_time = os.path.getmtime(file_path)
                    first_image_captured = True
                    print("Recorded first image time:", 
                          datetime.datetime.fromtimestamp(start_time))

                else:
                    if start_time is not None:
                        current_time = os.path.getmtime(file_path)
                        elapsed_seconds = current_time - start_time
                        print("Elapsed since first save:", int(elapsed_seconds), "seconds")

            except RuntimeError as e:
                print("Error capturing image:", e)

            # Delay between multiple gains in same cycle
            time.sleep(SHORT_INTERVAL)

        # End of cycle
        elapsed_time = time.time() - cycle_start_time
        remaining_time = CYCLE_INTERVAL - elapsed_time
        if remaining_time > 0:
            print(f"Cycle {cycle+1} finished in {elapsed_time:.2f}s. Sleeping {remaining_time:.2f}s.")
            time.sleep(remaining_time)
        else:
            print(f"Cycle {cycle+1} took {elapsed_time:.2f}s, exceeding CYCLE_INTERVAL. Moving on.")

        print("---")

    # Close camera after all cycles
    picam2.close()
    print("Timelapse complete. Camera closed.")

if __name__ == "__main__":
    main()
```
