## Supplementary File 3 for "ThermiQuant™ AquaStream: A portable instrument for quantitative colorimetric isothermal nucleic acid amplification reactions in paper and tube formats": AquaStream_software.py

```
#!/usr/bin/env python3

import sys
import os
import time
import datetime
import cv2
import queue
import json  # Added for JSON handling
from scipy.ndimage import gaussian_filter1d
from scipy.optimize import curve_fit
from scipy.special import wofz


from PyQt5.QtCore import QThread, pyqtSignal, QObject, Qt, QTimer
from PyQt5.QtWidgets import (
    QApplication, QWidget, QVBoxLayout, QHBoxLayout,
    QPushButton, QLabel, QPlainTextEdit, QTableWidget,
    QTableWidgetItem, QHeaderView, QMessageBox, QDialog,
    QLineEdit, QFormLayout, QDialogButtonBox, QComboBox  # Added QComboBox
)
from PyQt5.QtGui import QPixmap, QImage, QFont, QTextCursor, QColor

import pyqtgraph as pg
import RPi.GPIO as GPIO

# YOLO / patch / hue libraries
import numpy as np
import csv
import pandas as pd

import multiprocessing as mp

###############################################################################
# GLOBAL SETTINGS
###############################################################################
os.environ["QT_QPA_PLATFORM_PLUGIN_PATH"] = "/usr/lib/x86_64-linux-gnu/qt5/plugins"

APP_NAME        = "Paper LAMP Diagnostics"
ASPECT_RATIO    = 1.4
DISPLAY_SIZE    = (int(320*ASPECT_RATIO), 320)   # For GUI display
CAPTURE_SIZE    = (int(1920*ASPECT_RATIO), 1920) # Actual capture resolution

###############################################################################
# CREATE FOLDERS
###############################################################################
def make_new_folder(prefix):
    """Create a date-time-stamped folder with prefix on Desktop/Raw_Data, e.g., ASFV_1_20250101_161804."""
    timestamp_str = datetime.datetime.now().strftime("%Y%m%d_%H%M%S")
    desktop_path = os.path.join(os.path.expanduser("~"), 'Desktop', 'Raw_Data')
    os.makedirs(desktop_path, exist_ok=True)
    folder_name = f"{prefix}_{timestamp_str}"
    folder_path = os.path.join(desktop_path, folder_name)
    os.makedirs(folder_path, exist_ok=True)
    print(f"Created main folder: {folder_path}")
    return folder_path

###############################################################################
# DEFERRED CAMERA INIT
###############################################################################
def init_camera(capture_size, exposure_time, gain):
    """
    Creates & configures Picamera2, starts it, does auto-focus, 
    then locks the lens position. AWB is disabled for color consistency.
    Called in on_start_capture().
    """
    from picamera2 import Picamera2
    cam = Picamera2()

    preview_config = cam.create_preview_configuration(
        controls={"AfMode": 1, "AfTrigger": 0},
        main={"size": capture_size},
        buffer_count=1
    )
    cam.configure(preview_config)
    cam.start()

    # Let the camera warm up for focus
    time.sleep(3)
    cam.set_controls({"AfTrigger": 1})
    time.sleep(2)

    metadata = cam.capture_metadata()
    lens_position = metadata.get("LensPosition")
    # Lock the lens
    cam.set_controls({"AfMode": 0, "LensPosition": lens_position})

    # Set manual exposure, gain, and disable AWB
    # so the camera won't do any more auto adjustments.
    cam.set_controls({
        "ExposureTime": exposure_time,
        "AnalogueGain": gain,
        "AwbEnable": 0  # disable auto white balance
    })

    # Extra warm-up for stable color
    time.sleep(2)
    print("Camera initialized, lens locked, AWB disabled.")
    return cam

###############################################################################
# HELPER FUNCTIONS FOR MULTIPROCESSING
###############################################################################
def mp_initialize_yolo(model_path):
    """
    Global initializer for each worker process in the multiprocessing pool.
    """
    global YOLO_MODEL
    from ultralytics import YOLO
    YOLO_MODEL = YOLO(model_path)

def mp_process_image(args):
    """
    Worker process function for heavy tasks: YOLO inference, patch filtering,
    cropping, hue measurement, etc.
    """
    import cv2
    import numpy as np
    from ultralytics import YOLO
    (file_path, elapsed_minutes, frame_index, has_patches, first_patches, 
     model_path, csv_path, combined_dir, processed_dir,
     yolo_threshold, cycle_count, left_shrink, right_shrink,
     top_shrink, bottom_shrink) = args

    global YOLO_MODEL

    img = cv2.imread(file_path)
    if img is None:
        return {
            "status": "error",
            "error_msg": f"Cannot load {file_path}. Skipping.",
            "frame_index": frame_index
        }

    # No rotation as per instruction (removed rotation logic)

    # If not has_patches => run YOLO on first image
    final_patches = None
    is_first_frame = (frame_index == 1)

    if not has_patches:
        results = YOLO_MODEL.predict(source=img, conf=yolo_threshold)
        bounding_boxes = results[0].boxes.xyxy.cpu().numpy() if results else []
        filtered_patches = filter_segmented_patches(
            bounding_boxes, left_shrink, right_shrink, top_shrink, bottom_shrink
        )
        sorted_patches = sort_and_label_patches(filtered_patches)
        final_patches  = sorted_patches
    else:
        final_patches = first_patches

    # If no patches detected, emit a 'no_objects' status
    if not final_patches:
        return {
            "status": "no_objects",
            "error_msg": "No object detected. Either Light is turned off or cartridge is not loaded.",
            "frame_index": frame_index
        }

    # Save labeled image only for first or last
    labeled_image_path = None
    if frame_index == 1 or frame_index == cycle_count:
        labeled_img = draw_patches_on_image(img, final_patches)
        labeled_filename = f"labeled_frame_{frame_index}.jpg"
        labeled_path = os.path.join(processed_dir, labeled_filename)
        cv2.imwrite(labeled_path, labeled_img)
        labeled_image_path = labeled_path

    # Crop => measure hue => CSV
    crop_and_store_patches_combined(
        img, final_patches, combined_dir, frame_index, processed_dir,
        save_labeled_first_image=is_first_frame
    )
    combined_fname = f"ProcessedImage_{frame_index}.png"
    combined_path  = os.path.join(combined_dir, combined_fname)
    combined_img   = cv2.imread(combined_path)

    hue_dict = calculate_hue_for_patches_in_image_combined(combined_img, final_patches)
    save_hue_results_to_csv(hue_dict, round(elapsed_minutes, 2), csv_path)

    return {
        "status":           "ok",
        "frame_index":      frame_index,
        "elapsed_minutes":  elapsed_minutes,
        "labeled_path":     labeled_image_path,
        "patches":          final_patches,
        "hue_dict":         hue_dict,
        "has_patches":      True
    }

###############################################################################
# PATCH & HUE HELPER FUNCTIONS
###############################################################################
def filter_segmented_patches(bounding_boxes, ls, rs, ts, bs):
    patches = []
    for i, bbox in enumerate(bounding_boxes):
        x, y, x2, y2 = map(int, bbox[:4])
        w = (x2 - x)
        h = (y2 - y)
        # Shrink bounding box
        x  += int(ls * w)
        x2 -= int(rs * w)
        y  += int(ts * h)
        y2 -= int(bs * h)
        patches.append({
            'index': i+1,
            'bounding_box': (x, y, x2, y2),
        })
    if not patches:
        return []

    # Keep only patches above 20% of the largest
    largest_patch = max(patches, key=lambda p: bb_area(p['bounding_box']))
    largest_area  = bb_area(largest_patch['bounding_box'])
    threshold_area = 0.2 * largest_area
    patches = [p for p in patches if bb_area(p['bounding_box']) >= threshold_area]

    # Filter out weird outliers by approximate size
    sorted_by_area = sorted(patches, key=lambda p: bb_area(p['bounding_box']))
    mid_p = sorted_by_area[len(sorted_by_area)//2]
    ref_len = (mid_p['bounding_box'][2] - mid_p['bounding_box'][0])

    filtered_p = []
    for p in sorted_by_area:
        x1, y1, x2, y2 = p['bounding_box']
        w = (x2 - x1)
        h = (y2 - y1)
        if (0.25 * ref_len <= w <= 1.75 * ref_len) and (0.25 * ref_len <= h <= 1.75 * ref_len):
            filtered_p.append(p)
    return filtered_p

def bb_area(bb):
    x1, y1, x2, y2 = bb
    return (x2 - x1) * (y2 - y1)

def sort_and_label_patches(patches):
    if not patches:
        return []
    alpha = list("ABCDEFGHIJKLMNOPQRSTUVWXYZ")
    result = []
    row_count = 0
    fp = patches.copy()

    while fp:
        p0 = min(fp, key=lambda p: (p['bounding_box'][0]**2 + p['bounding_box'][1]**2))
        x1, y1, x2, y2 = p0['bounding_box']
        h  = (y2 - y1)
        my = (y1 + y2) / 2

        row = []
        for p in fp:
            xA1, yA1, xA2, yA2 = p['bounding_box']
            cy = (yA1 + yA2) / 2
            if (my - h / 0.75 <= cy <= my + h / 0.75):
                row.append(p)

        row.sort(key=lambda r: r['bounding_box'][0])
        row_label = alpha[row_count]
        for i, rP in enumerate(row, start=1):
            rP['label'] = row_label + str(i)

        row_count += 1
        result.extend(row)
        for r in row:
            if r in fp:
                fp.remove(r)
    return result

def draw_patches_on_image(image, patches):
    out = image.copy()
    for p in patches:
        if 'label' not in p:
            continue
        x, y, x2, y2 = p['bounding_box']
        lb = p['label']
        cv2.rectangle(out, (x, y), (x2, y2), (0, 255, 0), 4)
        txt_x, txt_y = x, y - 10
        (tw, th), _ = cv2.getTextSize(lb, cv2.FONT_ITALIC, 2.75, 5)
        cv2.rectangle(
            out,
            (txt_x, txt_y - th - 5),
            (txt_x + tw, txt_y + 5),
            (255, 255, 255), thickness=cv2.FILLED
        )
        cv2.putText(
            out, lb,
            (txt_x, txt_y),
            cv2.FONT_ITALIC, 2.75,
            (0, 0, 0), 10
        )
    return out

def crop_and_store_patches_combined(image, patches, out_dir, idx, processed_dir, save_labeled_first_image=False):
    import cv2
    import numpy as np

    patch_imgs = []
    labels = []
    for p in patches:
        if 'bounding_box' not in p or 'label' not in p:
            continue
        x1, y1, x2, y2 = p['bounding_box']
        sub_img = image[y1:y2, x1:x2]
        if sub_img.size == 0:
            continue
        sub_resized = cv2.resize(sub_img, (64, 64))
        patch_imgs.append(sub_resized)
        labels.append(p['label'])

    if not labels:
        print(f"No valid patches for frame {idx}")
        return

    row_letters = list(dict.fromkeys([l[0] for l in labels]))
    col_nums    = list(dict.fromkeys([int(l[1:]) for l in labels]))
    ph, pw      = 64, 64
    rows        = len(row_letters)
    cols        = len(col_nums)
    out_img     = np.zeros((ph * rows, pw * cols, 3), dtype=np.uint8)

    row_map = {l: i for i, l in enumerate(row_letters)}
    col_map = {n: i for i, n in enumerate(col_nums)}
    positions = []

    for i, lb in enumerate(labels):
        r_letter = lb[0]
        c_num = int(lb[1:])
        if r_letter not in row_map or c_num not in col_map:
            continue
        r = row_map[r_letter]
        c = col_map[c_num]
        sy = r * ph
        sx = c * pw
        out_img[sy:sy + ph, sx:sx + pw] = patch_imgs[i]
        positions.append((lb, sx, sy))

    out_fname = f"ProcessedImage_{idx}.png"
    out_path  = os.path.join(out_dir, out_fname)
    cv2.imwrite(out_path, out_img)
    print(f"Combined patches saved => {out_fname}")

    if save_labeled_first_image:
        labeled = out_img.copy()
        for (lb, sx, sy) in positions:
            cv2.putText(
                labeled, lb, (sx + 5, sy + 20),
                cv2.FONT_HERSHEY_SIMPLEX, 0.6,
                (0, 255, 0), 2
            )
        labeled_path = os.path.join(processed_dir, "first_combined_image_with_labels.png")
        cv2.imwrite(labeled_path, labeled)
        print("Saved first_combined_image_with_labels.png")

def calculate_hue_for_patches_in_image_combined(comb_img, patches):
    import cv2
    import numpy as np
    if comb_img is None or not patches:
        return {}
    hue_dict = {}
    labels = [p['label'] for p in patches if 'label' in p]
    if not labels:
        return hue_dict

    row_letters = list(dict.fromkeys([l[0] for l in labels]))
    col_nums    = list(dict.fromkeys([int(l[1:]) for l in labels]))
    if not row_letters or not col_nums:
        return hue_dict

    row_map = {l: i for i, l in enumerate(row_letters)}
    col_map = {n: i for i, n in enumerate(col_nums)}
    ph, pw = 64, 64

    for p in patches:
        lb = p['label']
        r_letter = lb[0]
        c_num    = int(lb[1:])
        if r_letter not in row_map or c_num not in col_map:
            continue
        r = row_map[r_letter]
        c = col_map[c_num]
        sy = r * ph
        sx = c * pw
        sub = comb_img[sy:sy + ph, sx:sx + pw]
        hsv = cv2.cvtColor(sub, cv2.COLOR_BGR2HSV)
        hue_vals = hsv[:, :, 0].astype(np.int16).ravel()

        filtered = [h for h in hue_vals if h < 31 or h > 159]
        modded   = [(h - 179 if h > 159 else h) for h in filtered]

        if modded:
            avg_hue = np.round(np.mean(modded), 1)
            hue_dict[lb] = float(avg_hue)
    return hue_dict

def save_hue_results_to_csv(hue_results, time_min, csv_path):
    labels = list(hue_results.keys())
    header = ["Time_min"] + labels
    file_exists = os.path.isfile(csv_path)
    with open(csv_path, mode="a", newline="") as csvfile:
        writer = csv.writer(csvfile)
        if not file_exists:
            writer.writerow(header)
        row_data = [time_min] + [hue_results.get(l, None) for l in labels]
        writer.writerow(row_data)

def voigt_fit(x, amplitude, center, sigma, gamma):
    """
    Voigt peak fit for derivative curve that enforces f(0)=0 by subtraction.
    (Same strategy as the reference code.)
    """
    z  = ((x - center) + 1j * gamma) / (sigma * np.sqrt(2))
    z0 = ((0 - center) + 1j * gamma) / (sigma * np.sqrt(2))
    norm = (sigma * np.sqrt(2*np.pi))
    return amplitude * (np.real(wofz(z))/norm - np.real(wofz(z0))/norm)

###############################################################################
# CAPTURE THREAD
###############################################################################
class CaptureThread(QThread):
    update_image     = pyqtSignal(str, int, float)
    capture_complete = pyqtSignal()
    capture_error    = pyqtSignal(str)  # Added error signal

    def __init__(self, image_queue, folder_paths, parent=None):
        super().__init__(parent)
        self.image_queue = image_queue
        self.folder_paths = folder_paths  # Dictionary containing folder paths
        self._running    = True
        self.picam2      = None

    def run(self):
        """Main capture logic. We init the camera here, skip the first warm-up frame, then capture CYCLES frames."""
        try:
            self.picam2 = init_camera(
                capture_size=self.folder_paths['capture_size'],
                exposure_time=self.folder_paths['exposure_time'],
                gain=self.folder_paths['gain']
            )  # fresh camera init each time "Start Capture" is pressed
        except Exception as e:
            print(f"Failed to initialize camera: {e}")
            self.capture_error.emit("Error in Capturing Image, Check Camera Cable Connection, or Restart Computer")
            return

        start_time = time.time()

        # --- Skip one warm-up frame (not saved, not processed) ---
        try:
            self.picam2.capture_array()  # just discard it
            print("Skipped a warm-up frame...")
        except Exception as e:
            print(f"Failed to skip warm-up frame: {e}")
            self.capture_error.emit("Error in Capturing Image, Check Camera Cable Connection, or Restart Computer")
            return

        # --- Actual capturing loop ---
        for frame_index in range(1, self.folder_paths['cycles'] + 1):
            if not self._running:
                break

            cycle_start = time.time()
            try:
                file_name = f"Gain{self.folder_paths['gain']}_Exp{self.folder_paths['exposure_time']}_Frame{frame_index}.jpg"
                file_path = os.path.join(self.folder_paths['main_folder_path'], file_name)
                self.picam2.capture_file(file_path)

                elapsed_minutes = (time.time() - start_time) / 60.0
                print(f"Captured image: {file_path}")

                # Emit signal to UI
                self.update_image.emit(file_path, frame_index, elapsed_minutes)
                # Add to queue for processing
                self.image_queue.put((file_path, elapsed_minutes, frame_index))

            except Exception as e:
                print(f"CaptureThread error: {e}")
                self.capture_error.emit("Error in Capturing Image, Check Camera Cable Connection, or Restart Computer")
                break

            elapsed_cycle_time = time.time() - cycle_start
            if self.folder_paths['cycle_interval'] > elapsed_cycle_time:
                self.sleep_interruptible(self.folder_paths['cycle_interval'] - elapsed_cycle_time)

        # Done capturing
        self.capture_complete.emit()
        print("Capture complete (CaptureThread). Cleaning up camera...")

        if self.picam2:
            self.picam2.stop()
            self.picam2.close()
            self.picam2 = None
        print("Camera fully stopped & closed in CaptureThread.")

    def stop(self):
        self._running = False
        print("Capture thread stopping...")

    def sleep_interruptible(self, duration):
        end_time = time.time() + duration
        while time.time() < end_time:
            if not self._running:
                break
            time.sleep(0.1)

###############################################################################
# PROCESSING THREAD (QThread + multiprocessing Pool)
###############################################################################
class ProcessingThread(QThread):
    update_labeled_image = pyqtSignal(str, int)
    update_hue_data      = pyqtSignal(float, dict)
    new_patches          = pyqtSignal(list)
    processing_done      = pyqtSignal(dict)  # Modified to emit classification data
    processing_error     = pyqtSignal(str)    # Added error signal

    def __init__(self, image_queue, folder_paths, parent=None):
        super().__init__(parent)
        self.image_queue   = image_queue
        self.folder_paths  = folder_paths  # Dictionary containing folder paths
        self._running      = True
        self.first_patches = []
        self.has_patches   = False
        self.process_count = 0

    def run(self):
        pool_size = 2  # Adjust if needed
        try:
            with mp.Pool(
                processes=pool_size,
                initializer=mp_initialize_yolo,
                initargs=(self.folder_paths['model_path'],)
            ) as pool:
                print("Multiprocessing pool started with size:", pool_size)

                outstanding_tasks = []
                while self._running:
                    try:
                        file_path, elapsed_minutes, frame_index = self.image_queue.get(timeout=0.5)
                        self.process_count += 1

                        args = (
                            file_path,
                            elapsed_minutes,
                            frame_index,
                            self.has_patches,
                            self.first_patches,
                            self.folder_paths['model_path'],
                            self.folder_paths['csv_filepath'],
                            self.folder_paths['combined_patches_dir'],
                            self.folder_paths['processed_data_dir'],
                            self.folder_paths['yolo_threshold'],
                            self.folder_paths['cycles'],
                            self.folder_paths['left_shrink'],
                            self.folder_paths['right_shrink'],
                            self.folder_paths['top_shrink'],
                            self.folder_paths['bottom_shrink']
                        )

                        result_async = pool.apply_async(
                            mp_process_image,
                            args=(args,),
                            callback=self.handle_result
                        )
                        outstanding_tasks.append(result_async)

                    except queue.Empty:
                        # If we have already processed CYCLES frames, we can break
                        if self.process_count >= self.folder_paths['cycles']:
                            all_done = all(r.ready() for r in outstanding_tasks)
                            if all_done:
                                break
                        continue
                    except Exception as e:
                        print(f"ProcessingThread exception: {e}")
                        self.processing_error.emit(f"Processing error: {e}")
                        continue

                # Wait for tasks if still running
                for r in outstanding_tasks:
                    if not r.ready():
                        try:
                            r.wait(timeout=10)
                        except:
                            pass

        except Exception as e:
            print(f"ProcessingThread failed to initialize pool: {e}")
            self.processing_error.emit(f"Processing pool initialization failed: {e}")
            return

        if self._running:
            normalized_csv = os.path.join(self.folder_paths['processed_data_dir'], "csv_normalized_hue_mean.csv")
            self.normalize_and_save_csv(self.folder_paths['csv_filepath'], normalized_csv)
            classification_data = self.obtain_classification(normalized_csv, self.folder_paths['hue_threshold'], self.folder_paths['tq_cutoff'], self.folder_paths['time_cutoff'])
            # Save Tq data to CSV
            tq_output_path = os.path.join(self.folder_paths['processed_data_dir'], "Tq_values.csv")
            classification_data['tq_values'].to_csv(tq_output_path, index=False)
            print(f"Saved Tq values => {tq_output_path}")
            # Emit signal with classification data
            self.processing_done.emit(classification_data)
            print("Processing thread finished all tasks.")

    def stop(self):
        self._running = False
        print("Processing thread stopping...")

    def handle_result(self, result):
        if not self._running or not result:
            return
        if result["status"] == "error":
            print(result["error_msg"])
            self.processing_error.emit(result["error_msg"])
            return
        elif result["status"] == "no_objects":
            print(result["error_msg"])
            self.processing_error.emit(result["error_msg"])
            return

        frame_index     = result["frame_index"]
        elapsed_minutes = result["elapsed_minutes"]
        labeled_path    = result["labeled_path"]
        hue_dict        = result["hue_dict"]

        if not self.has_patches and result["has_patches"]:
            self.has_patches   = True
            self.first_patches = result["patches"]
            self.new_patches.emit(self.first_patches)

        if labeled_path:
            self.update_labeled_image.emit(labeled_path, frame_index)

        self.update_hue_data.emit(elapsed_minutes, hue_dict)

    # ---------------------- Post-Processing ----------------------
    def normalize_and_save_csv(self, src_csv, dst_csv):
        if not os.path.isfile(src_csv):
            print(f"No CSV found => {src_csv}")
            return
        df = pd.read_csv(src_csv)
        if 'Time_min' not in df.columns:
            return

        norm_df = df.copy()
        for col in df.columns[1:]:
            if df[col].dtype in (np.float64, np.int64):
                first_val = df[col].iloc[0]
                norm_df[col] = df[col] - first_val

        norm_df.to_csv(dst_csv, index=False)
        print(f"Normalized CSV => {dst_csv}")

    def obtain_classification(self, normalized_csv, hue_threshold, tq_cutoff, time_cutoff):
        """
        Classify samples based on the hue and Tq thresholds.
        A sample is 'Pos' if hue >= hue_threshold AND Tq <= tq_cutoff.
        Otherwise, it's 'Neg'.
        Returns a dictionary containing classification data.
        """
        import pandas as pd

        # Load the normalized data
        data = pd.read_csv(normalized_csv)

        # Filter data based on time cutoff
        filtered_data = data[(data['Time_min'] >= self.folder_paths['min_time_cutoff']) & (data['Time_min'] <= time_cutoff)].reset_index(drop=True)


        # Dynamically determine sample columns (excluding "Time_min")
        sample_columns = [col for col in filtered_data.columns if col != 'Time_min']

        # Initialize lists to store positive and negative cases
        positive_cases = []
        negative_cases = []

        # Initialize a list to hold the classification table
        classification_table = []

        for col in sample_columns:
            # Check if any hue value meets the hue_threshold
            hue_condition = (filtered_data[col] >= hue_threshold).any()

            # Tq via Voigt on first-derivative, then derivative of the fit 
            x = filtered_data['Time_min'].to_numpy(dtype=float)

            # Smooth the signal (keep your smoothing)
            moving_avg = filtered_data[col].rolling(window=10, min_periods=1, center=True).mean()
            smoothed_data = gaussian_filter1d(moving_avg.to_numpy(), sigma=2)
            smoothed_series = pd.Series(smoothed_data, index=moving_avg.index)

            # First derivative of the smoothed signal (use gradient to respect x spacing)
            dy = np.gradient(smoothed_series.to_numpy(dtype=float), x)

            tq_time = np.inf  # default = fail (won't pass tq_cutoff)
            try:
                bounds = ([0, np.min(x), 1e-6, 1e-6], [np.inf, np.max(x), np.inf, np.inf])
                p0 = [float(np.max(dy) - np.min(dy)), float(x[int(np.argmax(dy))]), 2.0, 1.0]

                popt, _ = curve_fit(voigt_fit, x, dy, p0=p0, bounds=bounds, maxfev=20000)
                fitted_voigt = voigt_fit(x, *popt)

                # Optional R^2 gate like the reference code 
                residuals = dy - fitted_voigt
                ss_res = float(np.sum(residuals**2))
                ss_tot = float(np.sum((dy - np.mean(dy))**2))
                r2_voigt = 1 - ss_res / ss_tot if ss_tot != 0 else 0

                if r2_voigt >= 0.75:
                    second_deriv = np.gradient(fitted_voigt, x)  # ~ y''(t)
                    tq_time = float(x[int(np.argmax(second_deriv))])
            except Exception:
                pass

            # Classification based on both conditions
            if hue_condition and tq_time <= tq_cutoff:
                classification = "Pos"
                positive_cases.append(col)
            else:
                classification = "Neg"
                negative_cases.append(col)

            classification_table.append({"Sample": col, "Classification": classification})

        # Calculate Tq values for positive cases
        tq_values = {}
        for col in positive_cases:
            x = filtered_data['Time_min'].to_numpy(dtype=float)

            moving_avg = filtered_data[col].rolling(window=10, min_periods=1, center=True).mean()
            smoothed_data = gaussian_filter1d(moving_avg.to_numpy(), sigma=2)
            smoothed_series = pd.Series(smoothed_data, index=moving_avg.index)

            dy = np.gradient(smoothed_series.to_numpy(dtype=float), x)

            tq_time = np.inf
            try:
                bounds = ([0, np.min(x), 1e-6, 1e-6], [np.inf, np.max(x), np.inf, np.inf])
                p0 = [float(np.max(dy) - np.min(dy)), float(x[int(np.argmax(dy))]), 2.0, 1.0]

                popt, _ = curve_fit(voigt_fit, x, dy, p0=p0, bounds=bounds, maxfev=20000)
                fitted_voigt = voigt_fit(x, *popt)

                residuals = dy - fitted_voigt
                ss_res = float(np.sum(residuals**2))
                ss_tot = float(np.sum((dy - np.mean(dy))**2))
                r2_voigt = 1 - ss_res / ss_tot if ss_tot != 0 else 0

                if r2_voigt >= 0.75:
                    second_deriv = np.gradient(fitted_voigt, x)
                    tq_time = float(x[int(np.argmax(second_deriv))])

            except Exception as e:
                print(f"Voigt fit failed for {col}: {e}")

            tq_values[col] = tq_time if np.isfinite(tq_time) else "Tq_failed"

            
        # Update Tq values for negative cases as 'N/A'
        for col in negative_cases:
            tq_values[col] = 'N/A'

        # Create a DataFrame for the final Tq value table
        final_tq_table_df = pd.DataFrame(tq_values.items(), columns=["Sample", "Tq"])

        # Package the classification data
        classification_data = {
            "classification_table": classification_table,
            "tq_values": final_tq_table_df,
        }

        return classification_data

###############################################################################
# STREAM REDIRECTION
###############################################################################
class EmittingStream(QObject):
    textWritten = pyqtSignal(str)
    def write(self, text):
        self.textWritten.emit(str(text))
    def flush(self):
        pass

###############################################################################
# SEND REPORT DIALOG
###############################################################################
class SendReportDialog(QDialog):
    def __init__(self, parent=None):
        super().__init__(parent)
        self.setWindowTitle("Send Report")
        self.setModal(True)
        self.resize(300, 100)

        # Layout
        layout = QVBoxLayout()
        self.setLayout(layout)

        form_layout = QFormLayout()
        layout.addLayout(form_layout)

        self.email_input = QLineEdit()
        self.email_input.setText("vermalab.test.com")
        form_layout.addRow("Email:", self.email_input)

        # Buttons
        buttons = QDialogButtonBox(QDialogButtonBox.Ok | QDialogButtonBox.Cancel)
        buttons.accepted.connect(self.accept)
        buttons.rejected.connect(self.reject)
        layout.addWidget(buttons)

    def get_email(self):
        return self.email_input.text().strip()

###############################################################################
# MAIN WINDOW
###############################################################################
class MainWindow(QWidget):
    def __init__(self):
        super().__init__()
        self.setWindowTitle(APP_NAME)
        # We'll show maximized to allow minimize/close
        self.showMaximized()

        # Redirect stdout => console box
        self.emitting_stream = EmittingStream()
        self.old_stdout = sys.stdout
        sys.stdout = self.emitting_stream
        self.emitting_stream.textWritten.connect(self.append_text)

        # Layout
        self.main_layout = QVBoxLayout()
        self.setLayout(self.main_layout)

        self.top_layout = QHBoxLayout()
        self.main_layout.addLayout(self.top_layout)

        # Left: image display
        self.image_layout = QVBoxLayout()
        self.label_image  = QLabel()
        self.label_image.setAlignment(Qt.AlignCenter)
        self.image_layout.addWidget(self.label_image)

        self.label_progress = QLabel("Captured 0/0 Duration: 0 min")
        prog_font = QFont()
        prog_font.setPointSize(10)
        prog_font.setBold(True)
        self.label_progress.setFont(prog_font)
        self.image_layout.addWidget(self.label_progress, alignment=Qt.AlignCenter)

        self.top_layout.addLayout(self.image_layout, stretch=1)

        # Right: Plot and Table
        self.plot_or_table_layout = QVBoxLayout()
        self.top_layout.addLayout(self.plot_or_table_layout, stretch=1)

        # Initialize Plot Widget
        self.camera_plot = pg.PlotWidget()
        self.camera_plot.setBackground('w')
        self.camera_plot.showGrid(x=False, y=False)
        self.camera_plot.setTitle("Color LAMP Analysis", color='black', size='16pt', bold=True)
        self.camera_plot.setLabel("bottom", "Time (min)", color='black', **{'font-size': '16pt'})
        self.camera_plot.setLabel("left", "Color Value (Δ Hue)", color='black', **{'font-size': '16pt'})
        self.camera_plot.setYRange(-5, 15)

        # **Remove Legend Initialization**
        # self.legend = self.camera_plot.addLegend(offset=(0,10))  # Remove this line
        # Removed all legend-related code

        self.plot_or_table_layout.addWidget(self.camera_plot)

        # Initialize Table Widget (hidden by default)
        self.report_table = QTableWidget()
        self.report_table.setColumnCount(0)  # Will set dynamically
        self.report_table.setRowCount(0)     # Will set dynamically
        self.report_table.horizontalHeader().setSectionResizeMode(QHeaderView.Stretch)
        self.report_table.setFont(QFont("Arial", 12))
        self.report_table.verticalHeader().setVisible(False)
        self.report_table.setEditTriggers(QTableWidget.NoEditTriggers)
        self.report_table.setSelectionBehavior(QTableWidget.SelectRows)
        self.report_table.setSelectionMode(QTableWidget.SingleSelection)
        self.report_table.setStyleSheet("QTableWidget {background-color: white; color: black;}")
        self.report_table.hide()  # Hide initially
        self.plot_or_table_layout.addWidget(self.report_table)

        self.patch_data_x = {}
        self.patch_data_y = {}
        self.initial_hue_values = {}

        # Buttons
        self.button_layout = QHBoxLayout()
        self.main_layout.addLayout(self.button_layout)

        # 1. Add Dropdown for Chip Selection
        self.chip_dropdown = QComboBox()
        self.chip_dropdown.setFont(QFont("Arial", 14))
        # Dynamically populate the dropdown from JSON
        # Will do this after loading the JSON

        self.button_layout.addWidget(self.chip_dropdown)  # Add to left of buttons

        self.start_button = QPushButton("Start Capture")
        self.show_button  = QPushButton("Show Report")
        self.send_button  = QPushButton("Send Report")
        self.stop_button  = QPushButton("Stop")

        btn_font = QFont()
        btn_font.setPointSize(14)
        btn_font.setBold(True)
        for b in (self.start_button, self.show_button, self.send_button, self.stop_button):
            b.setFont(btn_font)
            b.setFixedHeight(int(b.sizeHint().height() * 1.5))  # FixedHeight updated here

        self.button_layout.addWidget(self.start_button)
        self.button_layout.addWidget(self.show_button)
        self.button_layout.addWidget(self.send_button)
        self.button_layout.addWidget(self.stop_button)

        # Console
        self.console_box = QPlainTextEdit()
        self.console_box.setReadOnly(True)
        font = QFont()
        font.setPointSize(10)
        self.console_box.setFont(font)
        self.console_box.setFixedHeight(100)
        self.main_layout.addWidget(self.console_box)

        # Threads
        self.image_queue       = queue.Queue()
        self.capture_thread    = None
        self.processing_thread = None

        # Patches for live overlay
        self.first_patches = None

        # Classification Data
        self.classification_data = None  # To store classification results

        # Connect
        self.start_button.clicked.connect(self.on_start_capture)
        self.show_button.clicked.connect(self.on_show_report)
        self.send_button.clicked.connect(self.on_send_report)
        self.stop_button.clicked.connect(self.on_stop_button)

        # Initial states
        self.enable_button(self.start_button, True)
        self.enable_button(self.show_button,  False)
        self.enable_button(self.send_button,  False)
        self.enable_button(self.stop_button,  False)

        # To keep track of the current experiment's folder paths
        self.current_folder_paths = None

        # Load chip settings
        self.chip_settings = self.load_chip_settings()
        if not self.chip_settings:
            QMessageBox.critical(self, "Error", "Failed to load chip settings. Application will exit.")
            sys.exit(1)

        # Populate the chip dropdown menu
        self.populate_chip_dropdown()

        # Display placeholder image
        self.display_placeholder_image()

    def load_chip_settings(self):
        """Load chip-specific settings from chip_settings.json."""
        script_dir = os.path.dirname(os.path.abspath(__file__))
        settings_path = os.path.join(script_dir, "chip_settings.json")
        if not os.path.exists(settings_path):
            print(f"chip_settings.json not found at {settings_path}")
            return None
        try:
            with open(settings_path, 'r') as f:
                settings = json.load(f)
            print("Chip settings loaded successfully.")
            return settings
        except json.JSONDecodeError as e:
            print(f"Error decoding chip_settings.json: {e}")
            return None

    def populate_chip_dropdown(self):
        """Populate the chip dropdown menu from the JSON file."""
        chip_names = sorted(self.chip_settings.keys())
        self.chip_dropdown.addItems(chip_names)
        self.chip_dropdown.setCurrentIndex(0)  # Set default to first chip

    def enable_button(self, btn, enable: bool):
        if enable:
            btn.setEnabled(True)
            btn.setStyleSheet("""
                QPushButton {
                    background-color: green; 
                    color: white; 
                    opacity: 1.0;
                }
            """)
        else:
            btn.setEnabled(False)
            btn.setStyleSheet("""
                QPushButton {
                    background-color: gray; 
                    color: black; 
                    opacity: 0.5;
                }
            """)

    ###########################################################################
    # START / STOP / SHOW / SEND
    ###########################################################################
    def on_start_capture(self):
        print("Start Capture clicked. Re-initializing camera & threads...")

        # Get selected chip
        selected_chip = self.chip_dropdown.currentText()
        print(f"Selected Chip: {selected_chip}")

        # Get chip-specific settings
        chip_config = self.chip_settings.get(selected_chip)
        if not chip_config:
            print(f"No configuration found for chip: {selected_chip}")
            QMessageBox.critical(self, "Error", f"No configuration found for chip: {selected_chip}")
            return

        # Create a new folder for this experiment with chip prefix
        main_folder_path = make_new_folder(selected_chip)

        # Subfolders
        combined_patches_dir = os.path.join(main_folder_path, 'combined_patches')  # Kept lowercase
        processed_data_dir   = os.path.join(main_folder_path, 'processed_data')    # Kept lowercase
        os.makedirs(combined_patches_dir, exist_ok=True)
        os.makedirs(processed_data_dir,   exist_ok=True)

        # CSV paths
        csv_filename = "csv_hue_mean.csv"  # kept lowercase
        csv_filepath = os.path.join(processed_data_dir, csv_filename)

        summary_csv_filename = "summary_report.csv"  # kept lowercase
        summary_csv_filepath = os.path.join(processed_data_dir, summary_csv_filename)

        # Define folder paths as a dictionary, including chip-specific settings
        folder_paths = {
            'main_folder_path': main_folder_path,
            'combined_patches_dir': combined_patches_dir,
            'processed_data_dir': processed_data_dir,
            'csv_filepath': csv_filepath,
            'summary_csv_filepath': summary_csv_filepath,
            'model_path': os.path.join(os.path.dirname(os.path.abspath(__file__)), "tube_pad_combined_best.pt"),
            'capture_size': CAPTURE_SIZE,
            'exposure_time': chip_config['exposure_time'],
            'gain': chip_config['gain'],
            'yolo_threshold': chip_config['yolo_threshold'],
            'hue_threshold': chip_config['hue_threshold'],
            'tq_cutoff': chip_config['tq_cutoff'],
            'time_cutoff': chip_config['time_cutoff'],
            'min_time_cutoff': chip_config['min_time_cutoff'],
            'cycles': chip_config['cycles'],
            'cycle_interval': chip_config['cycle_interval'],
            'left_shrink': chip_config['left_shrink'],
            'right_shrink': chip_config['right_shrink'],
            'top_shrink': chip_config['top_shrink'],
            'bottom_shrink': chip_config['bottom_shrink']
        }

        # Store current folder paths
        self.current_folder_paths = folder_paths

        # Clear old data
        self.camera_plot.clear()
        self.patch_data_x.clear()
        self.patch_data_y.clear()
        self.initial_hue_values.clear()
        self.label_progress.setText(f"Captured 0/{folder_paths['cycles']} Duration: 0 min")
        self.first_patches = None
        self.classification_data = None  # Reset classification data

        # Ensure plot is visible and table is hidden
        self.camera_plot.show()
        self.report_table.hide()

        # Start ProcessingThread
        from_processing = ProcessingThread(self.image_queue, folder_paths)
        from_processing.update_labeled_image.connect(self.on_labeled_image_ready)
        from_processing.update_hue_data.connect(self.on_new_hue_data)
        from_processing.new_patches.connect(self.on_new_patches)
        from_processing.processing_done.connect(self.on_processing_done)  # Connect to modified signal
        from_processing.processing_error.connect(self.handle_processing_error)  # Connect error signal
        self.processing_thread = from_processing
        self.processing_thread.start()

        # Start CaptureThread
        from_capture = CaptureThread(self.image_queue, folder_paths)
        from_capture.update_image.connect(self.on_image_captured)
        from_capture.capture_complete.connect(self.on_capture_complete)
        from_capture.capture_error.connect(self.handle_capture_error)  # Connect error signal
        self.capture_thread = from_capture
        self.capture_thread.start()

        # Buttons
        self.enable_button(self.start_button, False)
        self.enable_button(self.stop_button,  True)
        self.enable_button(self.show_button,  False)
        self.enable_button(self.send_button,  False)

    def on_stop_button(self):
        """Stop everything & close the app with confirmation."""
        print("Stop button clicked.")
        reply = QMessageBox.question(
            self, 'Confirm Stop',
            "Are you sure you want to stop the experiment and close the application?",
            QMessageBox.Yes | QMessageBox.No, QMessageBox.No
        )

        if reply == QMessageBox.Yes:
            if self.current_folder_paths:
                screenshot_path = os.path.join(self.current_folder_paths['processed_data_dir'], "appScreenshot.png")
                if not os.path.exists(screenshot_path):
                    # Ensure that the GUI has updated before taking the screenshot
                    QTimer.singleShot(0, self.save_screenshot)
                else:
                    print("Screenshot already exists. Not saving again.")
            else:
                print("No processed_data_dir found for screenshot.")

            print("User confirmed to stop. Closing application.")
            self.close()  # Triggers closeEvent
        else:
            print("User canceled the stop action.")

    def on_show_report(self):
        print("Show Report clicked.")
        if self.classification_data:
            self.populate_report_table()
            self.report_table.show()
            self.camera_plot.hide()
            self.enable_button(self.start_button, True)
            self.enable_button(self.show_button, False)
            self.enable_button(self.send_button, True)

            # Ensure that the table is fully rendered before taking a screenshot
            QTimer.singleShot(100, self.save_screenshot)  # Slight delay to ensure rendering
        else:
            print("No classification data available.")

    def on_send_report(self):
        print("Send Report clicked.")
        if self.classification_data:
            # Open Send Report Dialog
            dialog = SendReportDialog(self)
            if dialog.exec_() == QDialog.Accepted:
                email = dialog.get_email()
                if not email:
                    email = "vermalab.test.com"
                print(f"This is a placeholder. No report will be sent to: {email}")  # Placeholder for actual sending
                QMessageBox.information(self, "Send Report", f"This is a placeholder. No report will be sent to {email} successfully!")
            else:
                print("Send Report canceled by user.")

            # Implement report sending functionality here
            # For example, emailing the report or uploading to a server
            # Placeholder: print statement
            print("his is a placeholder. No report will be sent to.")  # Placeholder

            # Reset the image to placeholder
            self.display_placeholder_image()

            # Update button states
            self.enable_button(self.start_button, True)
            self.enable_button(self.stop_button,  False)
            self.enable_button(self.show_button,  False)
            self.enable_button(self.send_button,  False)
        else:
            print("No classification data available.")

    ###########################################################################
    # CAPTURE THREAD SIGNALS
    ###########################################################################
    def on_image_captured(self, file_path, frame_index, elapsed_minutes):
        img = cv2.imread(file_path)
        if img is not None:
            if self.first_patches is not None:
                img = draw_patches_on_image(img, self.first_patches)

            preview = cv2.resize(img, DISPLAY_SIZE)
            rgb = cv2.cvtColor(preview, cv2.COLOR_BGR2RGB)
            h, w, ch = rgb.shape
            qt_img = QImage(rgb.data, w, h, ch * w, QImage.Format_RGB888)
            self.label_image.setPixmap(QPixmap.fromImage(qt_img))
            self.label_image.setAlignment(Qt.AlignCenter)
            self.label_image.setStyleSheet("")  # Reset any styles
        else:
            print(f"Failed to read {file_path} for preview")

        self.label_progress.setText(
            f"Captured {frame_index}/{self.current_folder_paths['cycles']} Duration: {elapsed_minutes:.1f} min"
        )

    def on_capture_complete(self):
        print("Capture complete. Waiting for final processing...")

    ###########################################################################
    # PROCESSING THREAD SIGNALS
    ###########################################################################
    def on_labeled_image_ready(self, labeled_image_path, frame_index):
        img = cv2.imread(labeled_image_path)
        if img is not None:
            preview = cv2.resize(img, DISPLAY_SIZE)
            rgb = cv2.cvtColor(preview, cv2.COLOR_BGR2RGB)
            h, w, ch = rgb.shape
            qt_img = QImage(rgb.data, w, h, ch * w, QImage.Format_RGB888)
            self.label_image.setPixmap(QPixmap.fromImage(qt_img))
            self.label_image.setAlignment(Qt.AlignCenter)
            self.label_image.setStyleSheet("")  # Reset any styles
            print(f"Displayed labeled image for frame {frame_index}")

    def on_new_patches(self, patches):
        print("Received bounding-box patches from first YOLO detection.")
        self.first_patches = patches

    def on_new_hue_data(self, elapsed_minutes, raw_hue_dict):
        print(f"New raw hue data at {elapsed_minutes:.2f} min: {raw_hue_dict}")
        # Update initial hue if needed
        for label, raw_val in raw_hue_dict.items():
            if label not in self.initial_hue_values:
                self.initial_hue_values[label] = raw_val

        # Convert to normalized
        norm_dict = {}
        for label, raw_val in raw_hue_dict.items():
            base_val = self.initial_hue_values[label]
            diff_val = round(raw_val - base_val, 2)
            norm_dict[label] = diff_val

        # Add to data arrays & redraw
        for label, val in norm_dict.items():
            if label not in self.patch_data_x:
                self.patch_data_x[label] = []
                self.patch_data_y[label] = []
            self.patch_data_x[label].append(elapsed_minutes)
            self.patch_data_y[label].append(val)

        self.redraw_plot()

    def redraw_plot(self):
        self.camera_plot.clear()
        self.camera_plot.setBackground('w')
        self.camera_plot.showGrid(x=False, y=False)
        self.camera_plot.setTitle("Color LAMP Analysis", color='black', size='16pt', bold=True)
        self.camera_plot.setLabel("bottom", "Time (min)", color='black', **{'font-size': '16pt'})
        self.camera_plot.setLabel("left", "Color Value (Δ Hue)", color='black', **{'font-size': '16pt'})
        self.camera_plot.setYRange(-5, 15)

        # Define an expanded color list for more distinct colors
        colors = [
            '#1f77b4', '#ff7f0e', '#2ca02c', '#d62728', '#9467bd',
            '#8c564b', '#e377c2', '#7f7f7f', '#bcbd22', '#17becf',
            '#393b79', '#637939', '#8c6d31', '#843c39', '#7b4173',
            '#5254a3', '#6b6ecf', '#9c9ede', '#c7c7c7', '#dbdb8d',
            '#ffbb78', '#98df8a', '#ff9896', '#c5b0d5',
            '#c49c94', '#f7b6d2', '#c7c7c7', '#c7c7c7', '#c7c7c7'  # Add more if needed
        ]
        idx_color = 0
        all_times = []

        for label in sorted(self.patch_data_x.keys()):
            xdata = self.patch_data_x[label]
            ydata = self.patch_data_y[label]

            if not xdata or not ydata:
                continue

            # Sort data points by elapsed_minutes to ensure increasing time sequence
            sorted_pairs = sorted(zip(xdata, ydata), key=lambda pair: pair[0])
            sorted_x, sorted_y = zip(*sorted_pairs)

            all_times.extend(sorted_x)

            color = colors[idx_color % len(colors)]
            pen = pg.mkPen(color=color, width=2)
            self.camera_plot.plot(
                sorted_x, sorted_y,
                pen=pen,
                # name=label  # Remove name as legend is removed
            )
            idx_color += 1

        if all_times and max(all_times) > self.current_folder_paths['time_cutoff']:
            tmax = max(all_times)
            self.camera_plot.setXRange(tmax - self.current_folder_paths['time_cutoff'], tmax)
        else:
            self.camera_plot.enableAutoRange(axis=pg.ViewBox.XAxis, enable=True)

    def on_processing_done(self, classification_data):
        print("Processing done. Enabling 'Show Report' button.")
        self.classification_data = classification_data  # Store classification data
        self.enable_button(self.show_button, True)
        self.enable_button(self.send_button, False)  # Ensure "Send Report" is disabled until "Show Report" is pressed

        # Create metadata JSON
        self.create_metadata_file()

    ###########################################################################
    # CLEANUP
    ###########################################################################
    def closeEvent(self, event):
        """
        Properly stop threads, free camera, cleanup GPIO, restore stdout, etc.
        """
        print("Closing application window...")
        if self.capture_thread and self.capture_thread.isRunning():
            self.capture_thread.stop()
            self.capture_thread.wait()

        if self.processing_thread and self.processing_thread.isRunning():
            self.processing_thread.stop()
            self.processing_thread.wait()

        GPIO.cleanup()
        sys.stdout = self.old_stdout
        event.accept()
        print("Application closed cleanly.")

    def append_text(self, text):
        self.console_box.moveCursor(QTextCursor.End)
        self.console_box.insertPlainText(text)

    ###########################################################################
    # REPORT TABLE POPULATION
    ###########################################################################
    def populate_report_table(self):
        """
        Populate the report_table based on classification_data.
        Format it as a grid where rows are letters (A, B, C) and columns are numbers (1, 2, 3, 4, 5).
        Each cell displays "A1 (Neg, N/A)" or "B2 (Pos, 15.2)".
        Also, save the summary report as a CSV file with columns: Label, Classification, Tq.
        """
        if not self.classification_data:
            print("No classification data to populate.")
            return

        classification_table = self.classification_data.get("classification_table", [])
        tq_values_df = self.classification_data.get("tq_values", pd.DataFrame())

        if tq_values_df.empty:
            print("No Tq values available.")
            return

        # Merge classification with Tq values
        merged_df = pd.merge(pd.DataFrame(classification_table), tq_values_df, on='Sample', how='left')

        # Extract row and column identifiers
        row_labels = sorted(set([sample[0] for sample in merged_df['Sample']]))
        col_labels = sorted(set([int(sample[1:]) for sample in merged_df['Sample'] if sample[1:].isdigit()]))

        num_rows = len(row_labels)
        num_cols = len(col_labels)

        self.report_table.clear()
        self.report_table.setRowCount(num_rows)
        self.report_table.setColumnCount(num_cols)

        # Set row and column headers
        self.report_table.setVerticalHeaderLabels(row_labels)
        self.report_table.setHorizontalHeaderLabels([str(c) for c in col_labels])

        # Initialize summary data for CSV
        summary_data = []

        # Populate the table
        for _, row in merged_df.iterrows():
            sample = row['Sample']
            classification = row['Classification']
            tq = row['Tq']

            row_id = sample[0]  # Letter part
            col_id = int(sample[1:])  # Number part

            try:
                row_idx = row_labels.index(row_id)
                col_idx = col_labels.index(col_id)
            except ValueError:
                # If the row or column label is not found, skip
                continue

            # Format the cell content
            cell_text = f"{sample} ({classification}, {tq})"

            table_item = QTableWidgetItem(cell_text)
            table_item.setFont(QFont("Arial", 12))
            table_item.setTextAlignment(Qt.AlignCenter)

            # Apply text styling for "Pos" classification
            if classification.lower() == "pos" or classification.lower() == "positive":
                table_item.setFont(QFont("Arial", 12, QFont.Bold))
                table_item.setForeground(QColor(0, 0, 255))  # Blue text

            self.report_table.setItem(row_idx, col_idx, table_item)

            # Add to summary data
            summary_data.append({
                "Label": sample,
                "Classification": classification,
                "Tq": tq
            })

        # Enhance table appearance
        self.report_table.setStyleSheet("""
            QTableWidget::item {
                border: 1px solid #d3d3d3;
            }
            QTableWidget::item:selected {
                background-color: #a8d0e6;
            }
            QHeaderView::section {
                background-color: #f0f0f0;
                padding: 4px;
                border: 1px solid #d3d3d3;
                font-weight: bold;
            }
        """)

        print("Report table populated with actual classification data.")

        # Save summary report as CSV
        if summary_data:
            summary_df = pd.DataFrame(summary_data)
            summary_df.to_csv(self.current_folder_paths['summary_csv_filepath'], index=False)
            print(f"Summary report saved to {self.current_folder_paths['summary_csv_filepath']}")
        else:
            print("No summary data to save.")

    ###########################################################################
    # SCREENSHOT FUNCTIONALITY
    ###########################################################################
    def save_screenshot(self):
        """
        Save a screenshot of the application window to the processed_data directory
        as 'appScreenshot.png' if it does not already exist.
        """
        if not self.current_folder_paths:
            print("No current experiment folder paths found. Cannot save screenshot.")
            return

        screenshot_path = os.path.join(self.current_folder_paths['processed_data_dir'], "appScreenshot.png")
        if not os.path.exists(screenshot_path):
            pixmap = self.grab()
            pixmap.save(screenshot_path)
            print(f"Screenshot saved to {screenshot_path}")
        else:
            print("Screenshot already exists. Not saving again.")

    ###########################################################################
    # CREATE METADATA JSON FILE
    ###########################################################################
    def create_metadata_file(self):
        """
        Create a metadata.json file in the processed_data directory containing
        all key information used in the experiment run.
        """
        if not self.current_folder_paths:
            print("No current experiment folder paths found. Cannot create metadata.")
            return

        metadata = {
            "chip_name": self.chip_dropdown.currentText(),
            "start_time": datetime.datetime.now().strftime("%Y-%m-%d %H:%M:%S"),
            "global_settings": {
                "cycles": self.current_folder_paths['cycles'],
                "cycle_interval": self.current_folder_paths['cycle_interval'],
            },
            "chip_specific_settings": {
                "exposure_time": self.current_folder_paths['exposure_time'],
                "gain": self.current_folder_paths['gain'],
                "yolo_threshold": self.current_folder_paths['yolo_threshold'],
                "hue_threshold": self.current_folder_paths['hue_threshold'],
                "tq_cutoff": self.current_folder_paths['tq_cutoff'],
                "time_cutoff": self.current_folder_paths['time_cutoff'],
                "min_time_cutoff": self.current_folder_paths['min_time_cutoff'],
                "left_shrink": self.current_folder_paths['left_shrink'],
                "right_shrink": self.current_folder_paths['right_shrink'],
                "top_shrink": self.current_folder_paths['top_shrink'],
                "bottom_shrink": self.current_folder_paths['bottom_shrink']
            },
            "model_path": self.current_folder_paths['model_path'],
            "capture_size": self.current_folder_paths['capture_size']
        }

        # Add any additional metadata as needed

        metadata_path = os.path.join(self.current_folder_paths['processed_data_dir'], "metadata.json")
        try:
            with open(metadata_path, 'w') as f:
                json.dump(metadata, f, indent=4)
            print(f"Metadata saved to {metadata_path}")
        except Exception as e:
            print(f"Failed to save metadata: {e}")

    ###########################################################################
    # ERROR HANDLING SIGNALS
    ###########################################################################
    def handle_capture_error(self, message):
        self.display_error_message(message)

        # Stop capture_thread if running
        if self.capture_thread and self.capture_thread.isRunning():
            self.capture_thread.stop()
            self.capture_thread.wait()

        # Stop processing_thread if running
        if self.processing_thread and self.processing_thread.isRunning():
            self.processing_thread.stop()
            self.processing_thread.wait()

        # Update button states
        self.enable_button(self.start_button, True)
        self.enable_button(self.stop_button,  False)
        self.enable_button(self.show_button,  False)
        self.enable_button(self.send_button,  False)

    def handle_processing_error(self, message):
        self.display_error_message(message)

        # Stop capture_thread if running
        if self.capture_thread and self.capture_thread.isRunning():
            self.capture_thread.stop()
            self.capture_thread.wait()

        # Stop processing_thread if running
        if self.processing_thread and self.processing_thread.isRunning():
            self.processing_thread.stop()
            self.processing_thread.wait()

        # Update button states
        self.enable_button(self.start_button, True)
        self.enable_button(self.stop_button,  False)
        self.enable_button(self.show_button,  False)
        self.enable_button(self.send_button,  False)

    ###########################################################################
    # IMAGE DISPLAY METHODS
    ###########################################################################
    def display_placeholder_image(self):
        """Display the placeholder image 'first_image.jpg'."""
        script_dir = os.path.dirname(os.path.abspath(__file__))
        first_image_path = os.path.join(script_dir, "first_image.jpg")
        if os.path.exists(first_image_path):
            img = cv2.imread(first_image_path)
            if img is not None:
                preview = cv2.resize(img, DISPLAY_SIZE)
                rgb = cv2.cvtColor(preview, cv2.COLOR_BGR2RGB)
                h, w, ch = rgb.shape
                qt_img = QImage(rgb.data, w, h, ch * w, QImage.Format_RGB888)
                self.label_image.setPixmap(QPixmap.fromImage(qt_img))
                self.label_image.setAlignment(Qt.AlignCenter)
                self.label_image.setStyleSheet("")  # Reset any styles
                print("Displayed placeholder image.")
            else:
                self.display_error_message("Failed to load placeholder image.")
        else:
            self.display_error_message("Placeholder image not found.")

    def display_error_message(self, message):
        """Display an error message in the image area."""
        self.label_image.setPixmap(QPixmap())  # Clear any existing pixmap
        self.label_image.setText(message)
        self.label_image.setAlignment(Qt.AlignCenter)
        self.label_image.setStyleSheet("QLabel { color : red; font-size: 16pt; }")

    ###########################################################################
    # CLEANUP
    ###########################################################################
    def closeEvent(self, event):
        """
        Properly stop threads, free camera, cleanup GPIO, restore stdout, etc.
        """
        print("Closing application window...")
        if self.capture_thread and self.capture_thread.isRunning():
            self.capture_thread.stop()
            self.capture_thread.wait()

        if self.processing_thread and self.processing_thread.isRunning():
            self.processing_thread.stop()
            self.processing_thread.wait()

        GPIO.cleanup()
        sys.stdout = self.old_stdout
        event.accept()
        print("Application closed cleanly.")

    def append_text(self, text):
        self.console_box.moveCursor(QTextCursor.End)
        self.console_box.insertPlainText(text)

    ###########################################################################
    # REPORT TABLE POPULATION
    ###########################################################################
    def populate_report_table(self):
        """
        Populate the report_table based on classification_data.
        Format it as a grid where rows are letters (A, B, C) and columns are numbers (1, 2, 3, 4, 5).
        Each cell displays "A1 (Neg, N/A)" or "B2 (Pos, 15.2)".
        Also, save the summary report as a CSV file with columns: Label, Classification, Tq.
        """
        if not self.classification_data:
            print("No classification data to populate.")
            return

        classification_table = self.classification_data.get("classification_table", [])
        tq_values_df = self.classification_data.get("tq_values", pd.DataFrame())

        if tq_values_df.empty:
            print("No Tq values available.")
            return

        # Merge classification with Tq values
        merged_df = pd.merge(pd.DataFrame(classification_table), tq_values_df, on='Sample', how='left')

        # Extract row and column identifiers
        row_labels = sorted(set([sample[0] for sample in merged_df['Sample']]))
        col_labels = sorted(set([int(sample[1:]) for sample in merged_df['Sample'] if sample[1:].isdigit()]))

        num_rows = len(row_labels)
        num_cols = len(col_labels)

        self.report_table.clear()
        self.report_table.setRowCount(num_rows)
        self.report_table.setColumnCount(num_cols)

        # Set row and column headers
        self.report_table.setVerticalHeaderLabels(row_labels)
        self.report_table.setHorizontalHeaderLabels([str(c) for c in col_labels])

        # Initialize summary data for CSV
        summary_data = []

        # Populate the table
        for _, row in merged_df.iterrows():
            sample = row['Sample']
            classification = row['Classification']
            tq = row['Tq']

            row_id = sample[0]  # Letter part
            col_id = int(sample[1:])  # Number part

            try:
                row_idx = row_labels.index(row_id)
                col_idx = col_labels.index(col_id)
            except ValueError:
                # If the row or column label is not found, skip
                continue

            # Format the cell content
            cell_text = f"{sample} ({classification}, {tq})"

            table_item = QTableWidgetItem(cell_text)
            table_item.setFont(QFont("Arial", 12))
            table_item.setTextAlignment(Qt.AlignCenter)

            # Apply text styling for "Pos" classification
            if classification.lower() == "pos" or classification.lower() == "positive":
                table_item.setFont(QFont("Arial", 12, QFont.Bold))
                table_item.setForeground(QColor(0, 0, 255))  # Blue text

            self.report_table.setItem(row_idx, col_idx, table_item)

            # Add to summary data
            summary_data.append({
                "Label": sample,
                "Classification": classification,
                "Tq": tq
            })

        # Enhance table appearance
        self.report_table.setStyleSheet("""
            QTableWidget::item {
                border: 1px solid #d3d3d3;
            }
            QTableWidget::item:selected {
                background-color: #a8d0e6;
            }
            QHeaderView::section {
                background-color: #f0f0f0;
                padding: 4px;
                border: 1px solid #d3d3d3;
                font-weight: bold;
            }
        """)

        print("Report table populated with actual classification data.")

        # Save summary report as CSV
        if summary_data:
            summary_df = pd.DataFrame(summary_data)
            summary_df.to_csv(self.current_folder_paths['summary_csv_filepath'], index=False)
            print(f"Summary report saved to {self.current_folder_paths['summary_csv_filepath']}")
        else:
            print("No summary data to save.")

    ###########################################################################
    # SCREENSHOT FUNCTIONALITY
    ###########################################################################
    def save_screenshot(self):
        """
        Save a screenshot of the application window to the processed_data directory
        as 'appScreenshot.png' if it does not already exist.
        """
        if not self.current_folder_paths:
            print("No current experiment folder paths found. Cannot save screenshot.")
            return

        screenshot_path = os.path.join(self.current_folder_paths['processed_data_dir'], "appScreenshot.png")
        if not os.path.exists(screenshot_path):
            pixmap = self.grab()
            pixmap.save(screenshot_path)
            print(f"Screenshot saved to {screenshot_path}")
        else:
            print("Screenshot already exists. Not saving again.")

    ###########################################################################
    # CREATE METADATA JSON FILE
    ###########################################################################
    def create_metadata_file(self):
        """
        Create a metadata.json file in the processed_data directory containing
        all key information used in the experiment run.
        """
        if not self.current_folder_paths:
            print("No current experiment folder paths found. Cannot create metadata.")
            return

        metadata = {
            "chip_name": self.chip_dropdown.currentText(),
            "start_time": datetime.datetime.now().strftime("%Y-%m-%d %H:%M:%S"),
            "global_settings": {
                "cycles": self.current_folder_paths['cycles'],
                "cycle_interval": self.current_folder_paths['cycle_interval'],
            },
            "chip_specific_settings": {
                "exposure_time": self.current_folder_paths['exposure_time'],
                "gain": self.current_folder_paths['gain'],
                "yolo_threshold": self.current_folder_paths['yolo_threshold'],
                "hue_threshold": self.current_folder_paths['hue_threshold'],
                "tq_cutoff": self.current_folder_paths['tq_cutoff'],
                "time_cutoff": self.current_folder_paths['time_cutoff'],
                "min_time_cutoff": self.current_folder_paths['min_time_cutoff'],
                "left_shrink": self.current_folder_paths['left_shrink'],
                "right_shrink": self.current_folder_paths['right_shrink'],
                "top_shrink": self.current_folder_paths['top_shrink'],
                "bottom_shrink": self.current_folder_paths['bottom_shrink']
            },
            "model_path": self.current_folder_paths['model_path'],
            "capture_size": self.current_folder_paths['capture_size']
        }

        # Add any additional metadata as needed

        metadata_path = os.path.join(self.current_folder_paths['processed_data_dir'], "metadata.json")
        try:
            with open(metadata_path, 'w') as f:
                json.dump(metadata, f, indent=4)
            print(f"Metadata saved to {metadata_path}")
        except Exception as e:
            print(f"Failed to save metadata: {e}")

    ###########################################################################
    # ERROR HANDLING SIGNALS
    ###########################################################################
    def handle_capture_error(self, message):
        self.display_error_message(message)

        # Stop capture_thread if running
        if self.capture_thread and self.capture_thread.isRunning():
            self.capture_thread.stop()
            self.capture_thread.wait()

        # Stop processing_thread if running
        if self.processing_thread and self.processing_thread.isRunning():
            self.processing_thread.stop()
            self.processing_thread.wait()

        # Update button states
        self.enable_button(self.start_button, True)
        self.enable_button(self.stop_button,  False)
        self.enable_button(self.show_button,  False)
        self.enable_button(self.send_button,  False)

    def handle_processing_error(self, message):
        self.display_error_message(message)

        # Stop capture_thread if running
        if self.capture_thread and self.capture_thread.isRunning():
            self.capture_thread.stop()
            self.capture_thread.wait()

        # Stop processing_thread if running
        if self.processing_thread and self.processing_thread.isRunning():
            self.processing_thread.stop()
            self.processing_thread.wait()

        # Update button states
        self.enable_button(self.start_button, True)
        self.enable_button(self.stop_button,  False)
        self.enable_button(self.show_button,  False)
        self.enable_button(self.send_button,  False)

    ###########################################################################
    # IMAGE DISPLAY METHODS
    ###########################################################################
    def display_placeholder_image(self):
        """Display the placeholder image 'first_image.jpg'."""
        script_dir = os.path.dirname(os.path.abspath(__file__))
        first_image_path = os.path.join(script_dir, "first_image.jpg")
        if os.path.exists(first_image_path):
            img = cv2.imread(first_image_path)
            if img is not None:
                preview = cv2.resize(img, DISPLAY_SIZE)
                rgb = cv2.cvtColor(preview, cv2.COLOR_BGR2RGB)
                h, w, ch = rgb.shape
                qt_img = QImage(rgb.data, w, h, ch * w, QImage.Format_RGB888)
                self.label_image.setPixmap(QPixmap.fromImage(qt_img))
                self.label_image.setAlignment(Qt.AlignCenter)
                self.label_image.setStyleSheet("")  # Reset any styles
                print("Displayed placeholder image.")
            else:
                self.display_error_message("Failed to load placeholder image.")
        else:
            self.display_error_message("Placeholder image not found.")

    def display_error_message(self, message):
        """Display an error message in the image area."""
        self.label_image.setPixmap(QPixmap())  # Clear any existing pixmap
        self.label_image.setText(message)
        self.label_image.setAlignment(Qt.AlignCenter)
        self.label_image.setStyleSheet("QLabel { color : red; font-size: 16pt; }")

    ###########################################################################
    # SEND REPORT DIALOG
    ###########################################################################
    def on_send_report(self):
        print("Send Report clicked.")
        if self.classification_data:
            # Open Send Report Dialog
            dialog = SendReportDialog(self)
            if dialog.exec_() == QDialog.Accepted:
                email = dialog.get_email()
                if not email:
                    email = "vermalab.test.com"
                print(f"No Report will be sent to: {email}")  # Placeholder for actual sending
                QMessageBox.information(self, "Send Report", f"No Report sent to {email} successfully!")
            else:
                print("Send Report canceled by user.")

            # Implement report sending functionality here
            # For example, emailing the report or uploading to a server
            # Placeholder: print statement
            print("Report sent successfully.")  # Placeholder

            # Reset the image to placeholder
            self.display_placeholder_image()

            # Update button states
            self.enable_button(self.start_button, True)
            self.enable_button(self.stop_button,  False)
            self.enable_button(self.show_button,  False)
            self.enable_button(self.send_button,  False)
        else:
            print("No classification data available.")

###############################################################################
def main():
    # If needed: mp.set_start_method("spawn") or "fork"
    # Uncomment the following line if you encounter issues with multiprocessing on your system
    # mp.set_start_method("spawn")
    app = QApplication(sys.argv)
    window = MainWindow()
    # We already call showMaximized() in the __init__
    window.show()
    sys.exit(app.exec_())

if __name__ == "__main__":
    main()
```
